# A remorin from *Nicotiana benthamiana* interacts with the *Pseudomonas* type-III effector protein HopZ1a and is phosphorylated by the immune-related kinase PBS1

**DOI:** 10.1101/409235

**Authors:** Philip Albers, Suayib Üstün, Katja Witzel, Max Kraner, Frederik Börnke

## Abstract

The plasma membrane is at the interface of plant-pathogen interactions and thus many bacterial type-III effector proteins (T3Es) target membrane-associated processes to interfere with immunity. The *Pseudomonas syringae* T3E is a host cell plasma membrane (PM)-localized effector protein that has several immunity associated host targets but also activates effector triggered immunity (ETI) in resistant backgrounds. Although HopZ1a has been shown to interfere with early defense signaling at the PM, no dedicated plasma membrane-associated HopZ1a target protein has been identified until now. We show here, that HopZ1a interacts with the PM-associated remorin protein NbREM4 from *Nicotiana benthamiana* in several independent assays. NbREM4 re-localizes to membrane sub-domains after treatment with the bacterial elicitor flg22 and transient overexpression of NbREM4 in *N. benthamiana* induces the expression of a subset of defense related genes. We can further show that NbREM4 interacts with the immune-related receptor-like cytoplasmic kinase PBS1 and is phosphorylated by PBS1 on several residues *in vitro*. Thus, we conclude that NbREM4 is associated with early defense signaling at the PM. The possible relevance of the HopZ1a/NbREM4 interaction for HopZ1a virulence and avirulence functions is discussed.

## Introduction

Plants possess a sophisticated and multi-layered immune system that generally prevents infection by potential pathogens (Jones and Dangl, 2006; Dodds and Rathjen, 2010). This type of resistance is usually based on the recognition of conserved microbe-associated molecular patterns (MAMPs) by plant cell surface localized pattern recognition receptors (PRRs), which upon activation trigger a suite of defense responses collectively preventing ingress of invading pathogens and leading to so called pattern triggered immunity (PTI) (Macho and Zipfel, 2015). However, adapted pathogens have evolved virulence strategies to overcome plant defense and to cause disease in a given host species. The injection of type III effector (T3E) proteins into the host cell is an efficient mechanism employed by many Gram-negative bacterial pathogens to suppress plant immunity and to promote disease development (Büttner, 2016; Khan et al., 2018a). T3Es have diverse biochemical activities to interfere with host cellular processes including proteases, acetyltransferases, E3-ubiquitin ligases, phosphatases and protein kinases to name just a few. Host targets of T3Es include signaling proteins, transcriptional regulators, the protein processing machinery as well as metabolic enzymes with a majority of targets involved in plant immunity (Büttner, 2016; Khan et al., 2018a). Typically, a suite of 20 – 40 T3Es are translocated from a given bacterium into the host cell that collectively dampen the PTI response below a threshold allowing for bacterial multiplication and disease progression during so called effector triggered susceptibility (ETS) (Jones and Dangl, 2006).

As a response, plants have evolved the ability to either directly or indirectly recognize specific effector proteins through resistance proteins, a class of intracellular receptor proteins that typically contain nucleotide-binding domains (NB) and leucine rich repeats (LRRs). Effector recognition by NB-LRR proteins results in an accelerated and amplified PTI response and in most cases leads to localized cell death at the site of infection termed hypersensitive response (HR) and eventually inducing effector-triggered immunity (Dodds and Rathjen, 2010; Cesari, 2018).

The T3E HopZ1a from *Pseudomonas syringae* pv *syringae* A2 strain is a member of the widely distributed YopJ superfamily of cysteine proteases/acetyltransferases produced by both plant and animal bacterial pathogens (Lewis et al., 2011). It is a low specificity T3E that has several distinct molecular targets in different plant species (Khan et al., 2018a). HopZ1a possess acetyltransferase activity and has been shown to acetylate tubulin which disrupts the plant cytoskeletal network, resulting in breakdown of cellular trafficking (Lee et al., 2012a). In soybean, HopZ1a has been demonstrated to interact with the isoflavone biosynthesis enzyme 2-hydroxyisoflavanone dehydratase (GmHID1). Binding of HopZ1a promotes GmHID1 degradation and subsequently increases susceptibility of soybean to *P. syringae* by decreasing the capacity of the host to synthesize the phytoalexin daidzein (Zhou et al., 2011). Other host targets of HopZ1a include jasmonate ZIM-domain (JAZ) proteins, which negatively regulate the expression of jasmonic acid (JA)-responsive genes. Acetylation of JAZ proteins by HopZ1a mediates their degradation and thus activates JA signaling (Jiang et al., 2013). Activation of JA-signaling antagonizes salicylic acid (SA)-mediated defense responses which are required for immunity against hemibiotrophic pathogens, such *P. syringae* (Zheng et al., 2012).

Transgenic expression of HopZ1a in Arabidopsis has been shown to suppress several outputs of PTI such as the production of reactive oxygen species (ROS) and the activation of MAP-kinase signaling (Lewis et al., 2014). These effects are not well explained by the current known HopZ1a target proteins and thus additional targets might exist.

HopZ1a has been shown to trigger an HR in *Arabidopsis thaliana* accession Columbia-0 (Col-0), rice, certain soybean genotypes, and *Nicotiana benthamiana* (Ma et al., 2006). In Arabidopsis, HopZ1a recognition depends on the NB-LRR protein ZAR1 (HOPZ-ACTIVATED RESISTANCE1) and the ZED1 (HOPZ-ETI-DEFICIENT1) pseudokinase (Lewis et al., 2010; Lewis et al., 2013). ZED1 itself does not possess kinase activity and the current model suggests that it interacts with ZAR1 and HopZ1a and is acetylated by HopZ1a, which is hypothesized to trigger the activation of ZAR1 (Lewis et al., 2013). Thus, ZED1 appears to be a decoy guarded by ZAR1 and senses the activity of HopZ1a in the plant cell. Recent evidence suggests that in Arabidopsis ZAR1 also recognizes the *Xanthomonas campestris* T3E AvrAC, requiring the ZED1-related kinase ZRK1 instead of ZED1, as well as the *Pseudomonas syringae* T3E HopF2a in a ZRK3-dependent manner (Wang et al., 2015; Seto et al., 2017). A ZAR1 orthologue was recently identified in *N. benthamiana* (Baudin et al., 2017); however, opposite to earlier observations (Ma et al., 2006), HopZ1a only triggered a ZAR1 dependent HR in *N. benthamiana* upon co-expression with Arabidopsis ZED1 (Baudin et al., 2017). Thus, it is currently unclear what the endogenous guardee of ZAR1 in *N. benthamiana* is and if this is a decoy to lure HopZ1a into the ZAR1 immune complex.

In this study, we identified a remorin protein NbREM4 as a novel interaction partner of HopZ1a in *N. benthamiana* and further studies suggest that NbREM4 interacts with the immune related RLCK PBS1. Biochemical studies suggest that PBS1 can specifically phosphorylate NbREM4 *in vitro* on several residues. A possible role of the PBS1/NbREM4 interaction module as a target for HopZ1a is discussed. We further show that HopZ1a triggers a ZAR1-dependent HR in *N. benthamiana* suggesting that a HopZ1a targets a thus far unknown guardee of ZAR1 in this plant species.

## Results

### HopZ1a interacts with a remorin from *Nicotiana benthamiana* in yeast

HopZ1a is known to have multiple target proteins in different plant species (Zhou et al., 2011; Lee et al., 2012b; Jiang et al., 2013). However, the target proteins known to date do not explain all HopZ1a– related phenotypes observed upon expression of the effector in plants (Lewis et al., 2014). Thus, HopZ1a is likely to possess additional, yet unidentified target proteins. In order to further understand HopZ1a function, we screened for proteins that interact with HopZ1a using a yeast two-hybrid (Y2H) cDNA library from tobacco (*Nicotiana tabacum*). This repeatedly identified different clones of a cDNA encoding a protein with high similarity to remorin-like proteins from different plant species (Genbank accession number XP_016479648.1), which we tentatively named NtRemorin. Remorins are plant-specific plasma membrane proteins, which may act as molecular scaffolds regulating signal transduction (Jarsch and Ott, 2011). One clone isolated during the screening that comprised the entire predicted NtRemorin ORF, except the start methionine, was used in a direct Y2H assay with HopZ1a. Reporter gene activation confirms the interaction between the two proteins in yeast (Figure 1) We sought to use *Nicotiana benthamiana* for further functional analysis of the HopZ1a/remorin interaction because of its amenability to molecular and cell biology manipulations (Goodin et al., 2008). A homology search identified a remorin-like protein with high similarity (90%) to NtRemorin encoded by the *N. benthamiana* genome (Niben101Scf00735g05005.1), tentatively dubbed NbRemorin (Supplementary Figure S1). A direct Y2H interaction assay suggests that also NbRemorin binds to HopZ1a in yeast and thus is suitable for a functional analysis of the interaction (Figure 1). In Arabidopsis, the remorin family consists of 16 members which fall into six different groups (Raffaele et al., 2007). When grouped into a phylogenetic tree of the Arabidopsis remorin protein family, NbRemorin most closely associates with group 4 remorins and thus will be referred to as NbREM4 from now on (Supplementary Figure S2). In order to investigate whether HopZ1a can also interact with other members of the *N. benthamiana* remorin family, two additional remorin isoforms, one from group 1 and one from group 6, were tested for their ability to bind HopZ1a in yeast. A direct Y2H assay suggests no interaction of HopZ1a with NbRem1 or NbRem6 (Supplementary Figure S3). Thus, although binding of HopZ1a to other remorins not tested in this experiment cannot be excluded, there appears to be at least some specificity of HopZ1a to interact with NbREM4.

**Figure 1.**
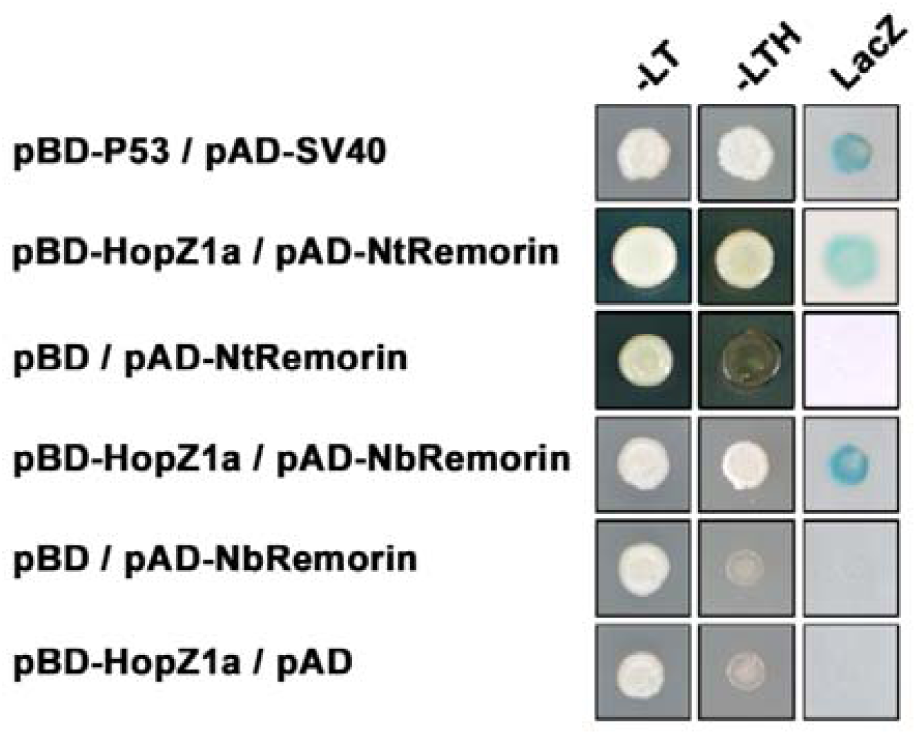
HopZ1a interacts with a remorin in yeast two-hybrid assays. HopZ1a fused to the GAL4 DNA binding domain (BD) was expressed in combination with remorin fused to the GAL4 activation domain (AD) in yeast strain Y190. Cells were grown on selective media before a LacZ filter assay was performed. pSV40/p53 served as positive control while the empty AD vector served as negative control. NtRemorin, *N. tabacum* remorin; NbRemorin, *N. benthamiana* remorin. – LT, yeast growth on medium without Leu and Trp. – HLT, yeast growth on medium lacking His, Leu, and Trp, indicating expression of the *HIS3* reporter gene. LacZ, activity of the *lacZ* reporter gene.

The NbREM4 polypeptide comprises 296 amino acids and has a predicted molecular weight of 33 kDa. It features the typical domain structure found in remorins from different plant species (Raffale et al., 2007) with a conserved C-terminal signature region (Pfam domain Remorin_C; PF03763) that encodes a predicted coiled-coil motif and a putative membrane-anchoring motif and a variable N-terminal part (Figure S1). In order to map the region of HopZ1a binding to NbREM4, we co-transformed constructs comprising the NbREM4 N-terminal region or the C-terminal region of the protein together with HopZ1a in a yeast reporter strain. As shown in Figure 2, the conserved NbREM4 C-terminus is necessary and sufficient for the interaction with HopZ1a. However, the apparently weaker interaction indicates that the N-terminal part of the protein likely also contributes to HopZ1a binding.

**Figure 2.**
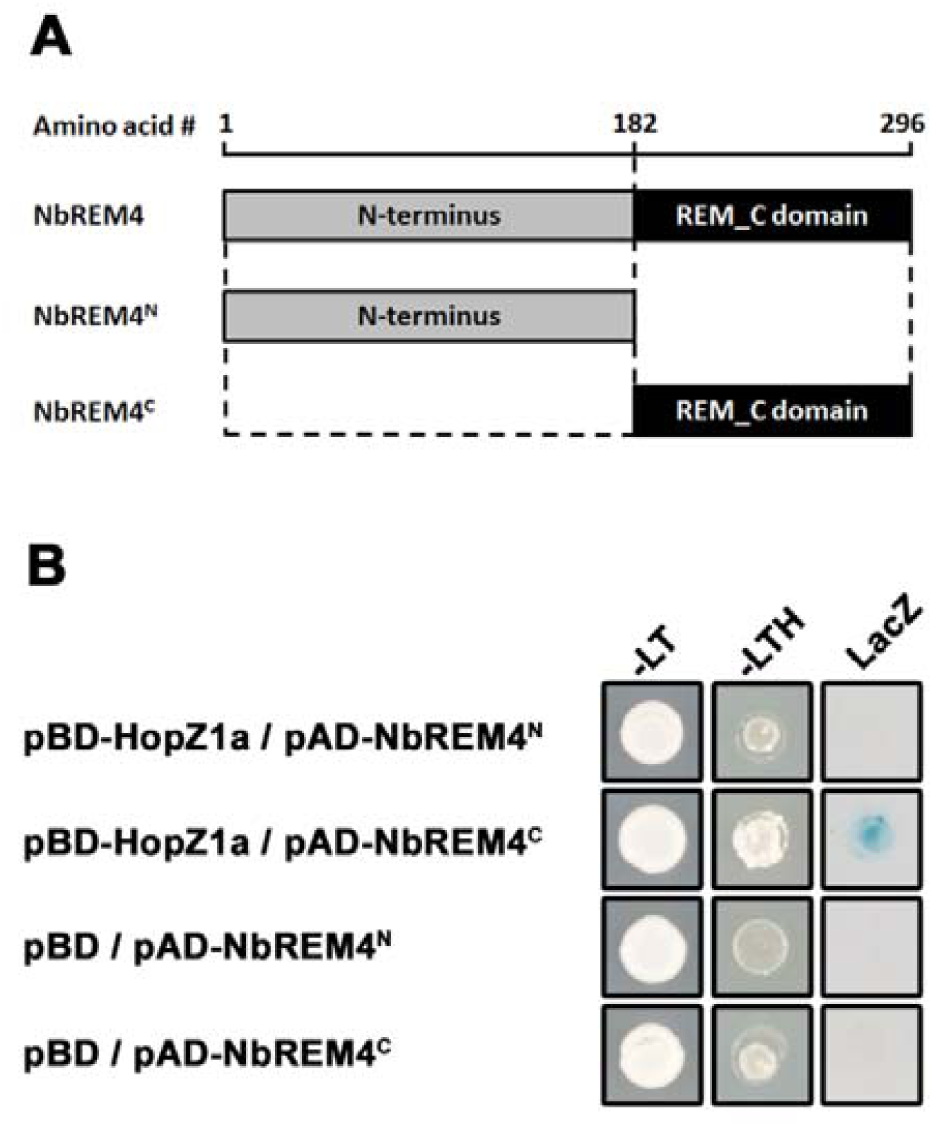
The NbREM4 C-terminus is required and sufficient for HopZ1a interaction in yeast. **A,** schmematical representation of the NbREM4 domain structure. **B,** HopZ1a interacts with the NbREM4 C-terminal region (NbREM4^C^) comprising amino acids 182 – 296 but not with the N-terminus of NbREM4 (NbREM^N^) covering amino acids 1 – 181. Cells were grown on selective media before a LacZ filter assay was performed. pSV40/p53 served as positive control while the empty BD vector served as negative control. – LT, yeast growth on medium without Leu and Trp. – HLT, yeast growth on medium lacking His, Leu, and Trp, indicating expression of the *HIS3* reporter gene. LacZ, activity of the *lacZ* reporter gene.

Remorins have been reported to form hetero and homo-oligomers and thus we tested the ability of NbREM4 to self-interact or to bind to other remorin isoforms. A direct Y2H assay indicates that indeed NbREM4 is able to form homomers in yeast but does not bind to two other remorin isoforms tested (Supplementary Figure S4). To confirm this interaction in planta, we performed BiFC using *N. benthamiana* leaves, where a robust fluorescence signal indicated oligomerization of NbREM4 at the PM of infiltrated tobacco epidermal cells (Supplementary Figure S4).

### HopZ1a interacts with NbREM4 *in planta* and in vitro

Next, the subcellular localization of both protein partners *in planta* was examined to determine whether co-localization as a prerequisite of *in vivo* interaction can be observed. The green fluorescent protein (GFP) was fused to the C-terminus of HopZ1^C/A^ carrying a cysteine to alanine substitution at position C216 within the conserved catalytic triad of the effector, and the fusion protein was expressed in leaves of *N. benthamiana* using *Agrobacterium*-infiltration. The HopZ1a^C/A^ variant was used for the experiments to prevent elicitation of a hypersensitive response in *N. benthamiana* that has been shown to depend on its catalytic activity (Ma et al., 2006). In accordance with previous findings and the presence of a myristoylation motif at glycine 2 of the HopZ1a (Lewis et al., 2008), HopZ1a^C/A^-GFP expression generated a fluorescence signal in the periphery of the cell, indicating plasma membrane localization of the fusion protein (Figure 3A). In a similar approach NbREM4 was tagged at the N-terminus with GFP and transient expression of the GFP-NtRem4 protein in *N. benthamiana* yielded a fluorescence signal resembling that of a PM associated protein (Figure 3B). To further corroborate this finding, membranes of GFP-NbREM4 expressing cells were labelled with the fluorescence dye FM4-64 and microscopic analysis revealed substantial overlap between the GFP and the FM4-64 fluorescence signal, confirming PM localization of NbREM4 (Figure 3B). Thus, both HopZ1a and NbREM4 localize to the plant cell PM and therefore could interact *in vivo*.

**Figure 3.**
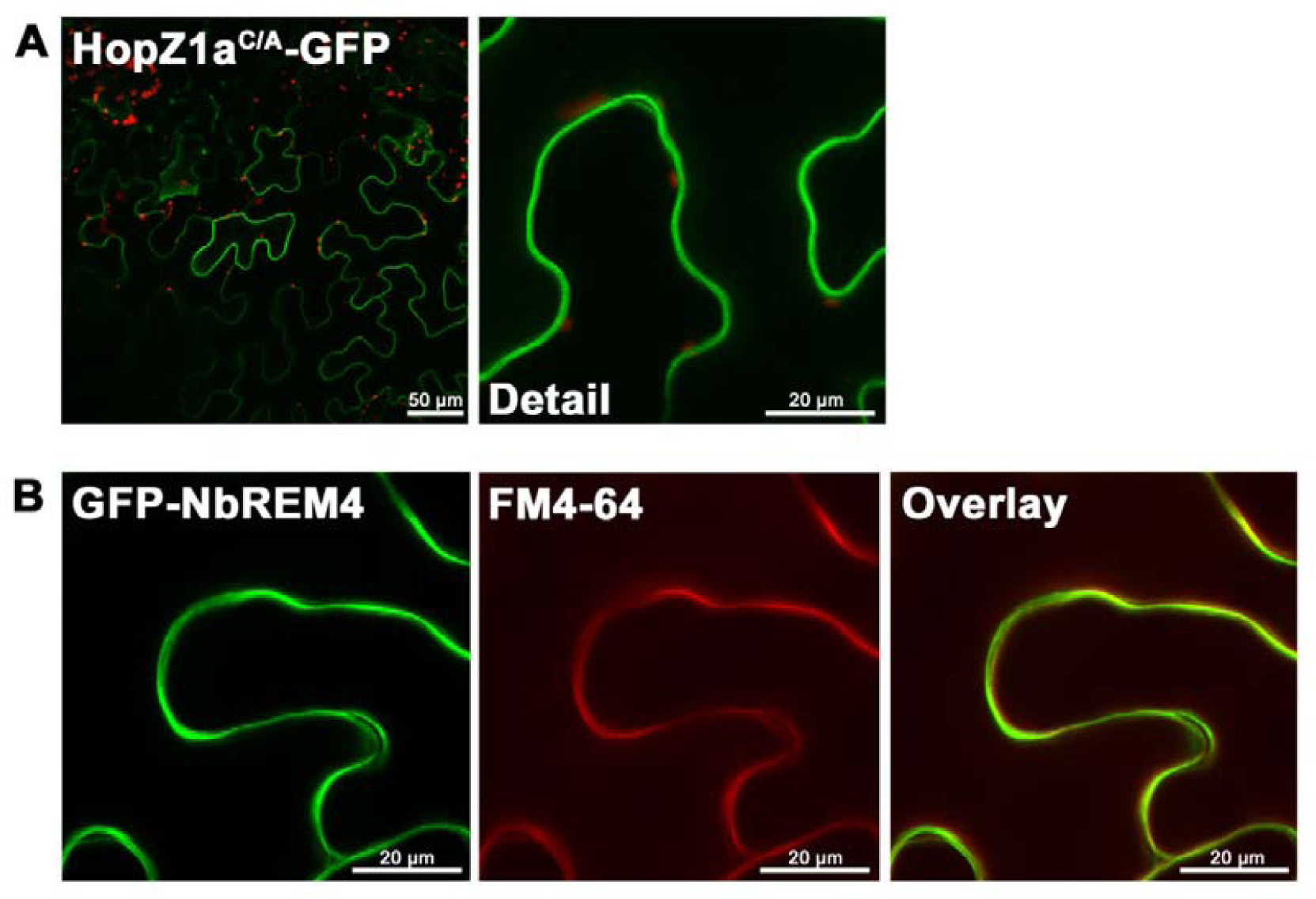
Subcellular localization of HopZ1a and NbREM4 *in planta*. **A,** Subcellular localization of HopZ1a^C/A-^GFP in *N. benthamiana* leaves transiently transformed by *Agrobacterium*-infiltration. For confocal laser scanning microscopy samples were taken 48 hpi. **B,** Subcellular localization of GFP-NbREM4. Membranes were stained with FM4-64 (middle) and green and red fluorescence channels were recorded separately to prevent bleed through. The resulting fluorescence images were merged (right). Pictures were taken 48 hpi.

To verify the interaction of HopZ1a and NbREM4 *in planta*, bimolecular fluorescence complementation (BiFC) assays were performed in *N. benthamiana* using transient expression via *Agrobacteria*. Strong YFP fluorescence was observed when a combination of HopZ1a-Venus^C^ with Venus^N^-NbREM4 was expressed, demonstrating that both proteins interact inside plant cells (Figure 4A). In accordance with PM localization of HopZ1a as well as of NbREM4, the YFP signal appeared to be confined to the PM as no fluorescence surrounding the chloroplasts could be detected which would be indicative for a cytosolic localization of the interaction (Figure 4A). Negative controls including unrelated proteins yielded no fluorescence signal, indicating the specificity of the interaction (Supplementary Figure S5). In addition, mutation of the myristoylation motif at glycine 2 of HopZ1a abolished the interaction with NbREM4 (Figure 4A), which is in accordance with a release of the HopZ1^G/A^ protein from the plasmamembrane (Lewis et al., 2008).

**Figure 4.**
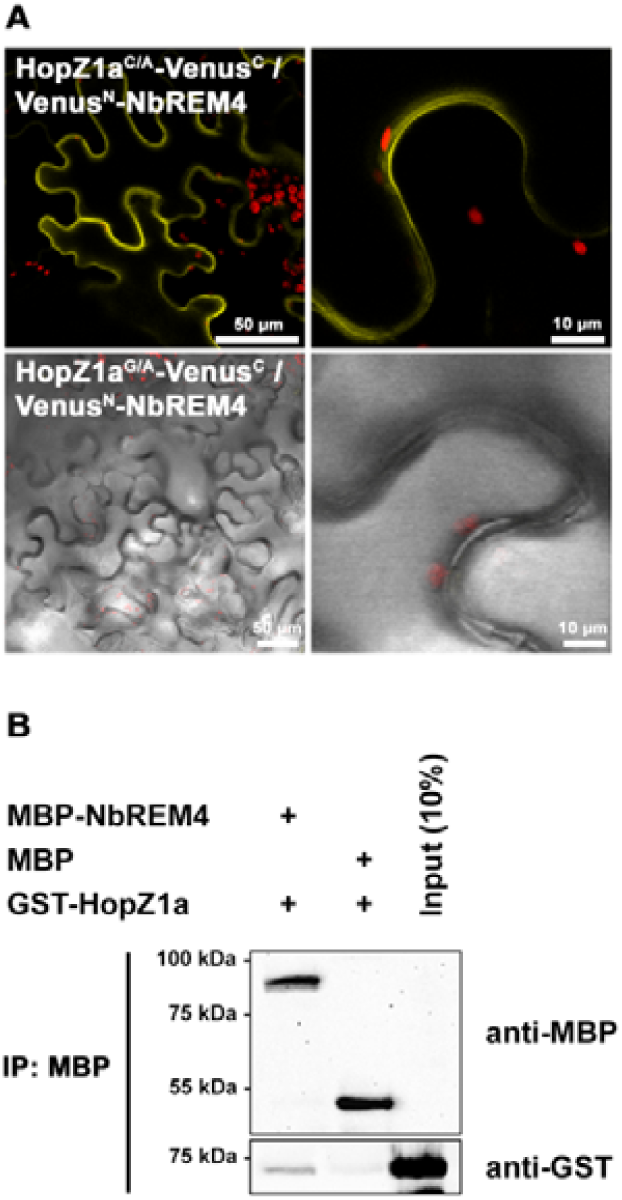
Interaction of HopZ1a with NbREM4 *in planta* and *in vitro*. **A,** BiFC in planta interaction studies of HopZ1a variants with NbREM4. YFP confocal microscopy images show *N. benthamiana* leaf epidermal cells transiently expressing HopZ1a^C/A^-Venus^C^ in combination with Venus^N^-NbREM4. A close-up of the same cells shows that the YFP fluorescence of HopZ1a^C/A^-Venus^C^/Venus^N^-NbREM4 aligns with the plasma membrane. The lower panel shows DIC images of leave epidermal cells expressing a combination of Venus^N^-NbREM4 and HopZ1a^G/A^-Venus^C^, indicating that a mutation at the myristoylation site at G2 of HopZ1a abolishes the interaction with NbREM4. **B,** *In vitro* pull-down assay showing physical interaction of HopZ1a with NbREM4. MBP-NbREM4 and GST-HopZ1a were expressed in *E. coli*. Pull-down was performed using amylose resin. Proteins were detected in an immunoblot using antibodies as indicated.

To exclude that the interaction between HopZ1a and NbREM4 is mediated by a third eukaryotic protein an *in vitro* pull-down assay was performed. To this end, recombinant glutathione S-transferase (GST) tagged HopZ1a was incubated with maltose-binding protein (MBP) tagged NbREM4. Protein complexes were pulled-down using amylose resin which binds MBP and precipitated proteins were subsequently detected using either anti-GST or anti-MBP antibodies. A western blot revealed that GST-HopZ1a was pulled down together with MBP-NbREM4, demonstrating a direct physical interaction of both proteins which does not require additional factors (Figure 4B). MBP alone was not able to pull down GST-HopZ1a indicating specificity of the *in vitro* interaction (Figure 4B).

Taken together, these data suggest that the *Pseudomonas* T3E HopZ1a directly interacts with the remorin NbREM4 at the PM of plant cells.

### Overexpression of NbREM4 affects defense gene expression

Remorins from different plant species have been associated with plant defense responses (Jarsch and Ott, 2011; Bozkurt et al., 2014; Son et al., 2014; Fu et al., 2018). Given the fact that the majority of *Pseudomonas* T3Es target immunity related functions inside the host cell (Büttner, 2016; Khan et al., 2018a), the interaction of NbREM4 with HopZ1a points towards an involvement of NbREM4 in some sort of plant defense response. In order to provide first insights into an *in planta* function of NbREM4 a GFP-tagged version of the protein under the control or the constitutive CaMV35S promoter was transiently expressed in leaves of *N. benthamiana* using *Agrobacterium*-infiltration. A phenotypic inspection of GFP-NbREM4 expressing leaves revealed that overexpression of the remorin protein led to the development of chloroses in infiltrated areas 3 dpi (days post infiltration) which progressed into tissue collapse until 6 dpi (Figure 5A). An anti-GFP western blot suggests that GFP-NbREM4 protein levels are highest 1 and 2 dpi and decline to almost undetectable levels until 6 dpi, which is in accordance with leaf phenotype development (Figure 5C). To quantitatively assess tissue damage upon GFP-NbREM4 expression, electrolyte leakage from infiltrated leaves was measured at different time points. Ion leakage was significantly increased in GFP-NbREM4 expressing leaves compared to leaves transformed with the empty vector from 3 dpi onwards (Figure 5B), indicating that GFP-NbREM4 expression leads to membrane damage. To monitor GFP-NbREM4 triggered molecular changes expression of selected defense related genes was measured 24 dpi and 48 hpi, before any phenotypic changes became visible. The genes *Pti5, Acre31* and *Gras2* have been demonstrated previously to be responsive to both a nonadapted bacterium and a semi-virulent bacterial pathogen in *N. benthamiana* and thus have been implicated in PTI (Nguyen et al., 2010). Expression of *Pti5* and *Acre31* was induced in GFP-NbREM4 expressing plants at 48 hpi (Figure 6). For *Gras2*, expression was higher at 24 hpi compared to 48hpi, whereas *Acre31* expression was maintained at similar levels between the two time points. Although not significantly upregulated, *Gras2* expression was increased by trend at 48 hpi in GFP-NbREM4 infiltrated leaves (Figure 6). In addition, GFP-NbREM4 overexpression led to the significant up-regulation of the pathogenesis related genes *Hin1* and *Hsr201* (Figure 6). Taken together, transient overexpression of NbREM4 in leaves of *N. benthamiana* induces expression of immunity related genes 48 hpi and leads to tissue collapse at later time points.

**Figure 5.**
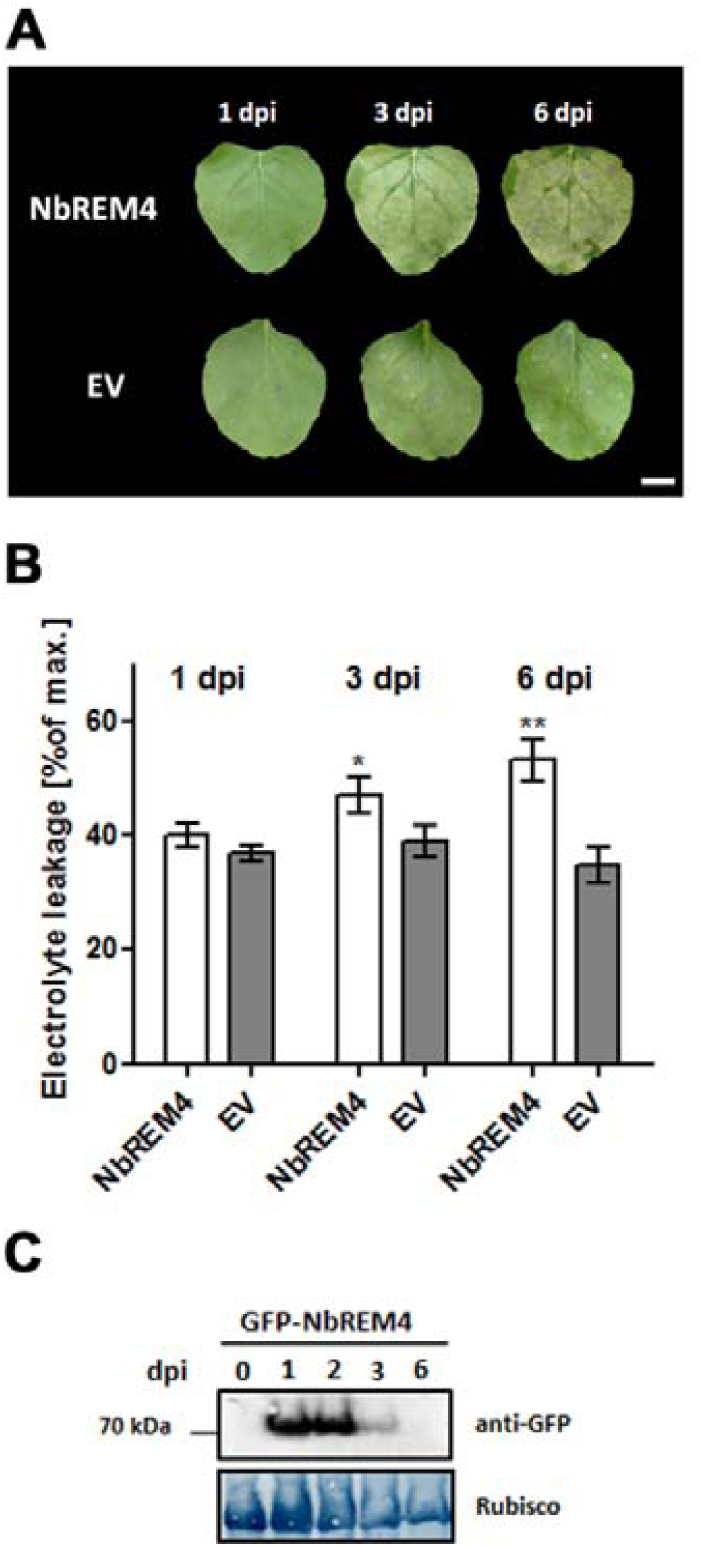
Transient over-expression of NbREM4 in leaves of *N. benthamiana* leads to tissue damage. **A,** Time course of phenotype development in *N. benthamiana* leaves infiltrated with *Agrobacterium tumefaciens* strains that mediate T-DNA-based transfer of NbREM4 and empty vector (EV). dpi = days post infiltration. B, Ion leakage was measured in plants transiently expressing NbREM4 and EV at time points indicated. Bars represent the average ion leakage measured for triplicates of six leaf disks each, and the error bars indicate SD. The asterisk indicates a significant difference (\**P* < 0.05, \*\**P* < 0.01) based on results of a Student’s *t*-test. The experiment has been repeated three times with similar results. **C,** Protein extracts from *N. benthamiana* leaves transiently expressing GFP-NbREM4 at the time points indicated were prepared. Equal volumes representing approximately equal protein amounts of each extract were immunoblotted and proteins were detected using anti-GFP antiserum. Amido black staining served as a loading control.

**Figure 6.**
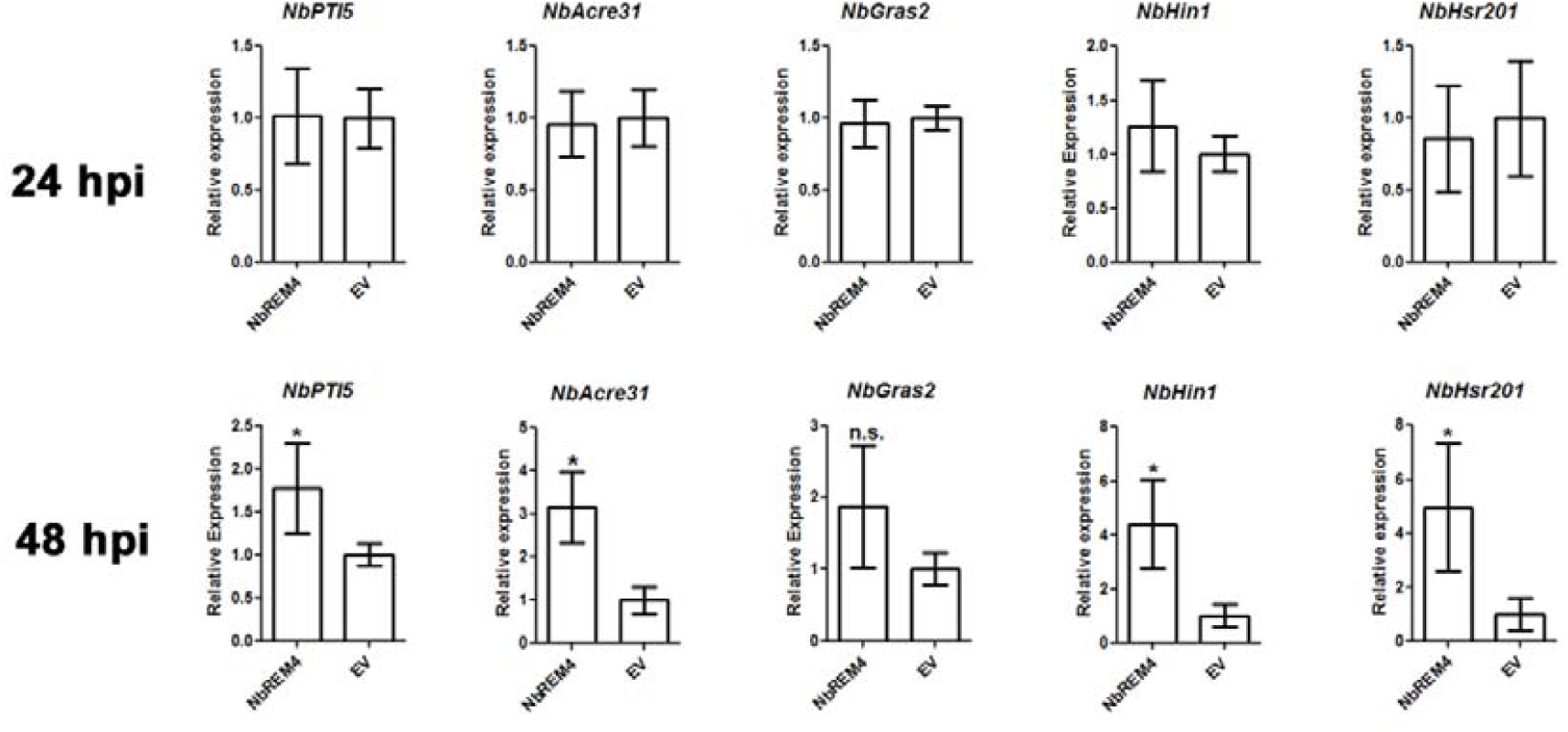
Expression of selected defense-related genes in *N. benthamiana* leaves transiently expressing NbREM4. Quantitative real-time PCR (RT-PCR) of indicated defense-related genes was carried out on samples taken from *N. benthamiana* leaves transiently expressing NbREM4 or the empty vector control (EV) at the time points indicated. *Actin* was used as a reference gene. Each bar represents the mean of four biological replicates ± SE. * marks significant differences (*P* > 0.05) according to Student’s *t*-test.

### NbREM4 interacts with the immune related kinase PBS1

It has been hypothesized that certain remorin isoforms play a dynamic role as scaffold proteins in plant innate immune signaling (Benschop et al., 2007; Jarsch and Ott, 2011). Thus, we sought to identify plant proteins capable of interacting with NbREM4 which could provide further information about NbREM4 function. To this end, a Y2H screening of a tobacco cDNA library using NbREM4 as a bait was conducted. This identified a PBS1-like protein as a potential NbREM4 interactor in yeast, which encodes a receptor-like cytoplasmic kinase (RLCK). PBS1-like proteins belong to subfamily VII of RLCKs that include the founding member avrPphB-susceptible 1 (PBS1) (Shao et al., 2003) and the Botrytis-induced kinase 1 (BIK1) (Laluk et al., 2011) as well as other PBS-like (PBL) proteins implicated in plant immune signaling (Lin et al., 2013; Rao et al., 2018).

We isolated a full-length sequence of the NbREM4 interacting PBS-like from *N. benthamiana* (Niben101Scf02086g00004.1). The protein comprises 390 amino acids and shares 71 % similarity with PBS1 from Arabidopsis (Supplementary Figure S6). Just as AtPBS1, the *N. benthamiana* PBS-like protein carries a putative myristoylation signal at its N-terminus which likely mediates PM association and contains a single catalytic kinase domain. Based on these analogies we termed the *N. benthamiana* PBS-like protein NbPBS1. A direct Y2H assay revealed that full-length NbPBS1 as well as Arabidopsis (At)PBS1 is capable of interacting with NbREM4 in yeast (Figure 7A). In order to investigate whether NbREM4 specifically interacts with PBS1 or also with other member of the PBL family we tested binding of the remorin to other RLCKs from Arabidopsis. As shown in Figure S7, NbREM4 interacted with AtPBS1 but not with any other RLCK tested, indicating a certain degree of specificity in the interaction of both proteins.

**Figure 7.**
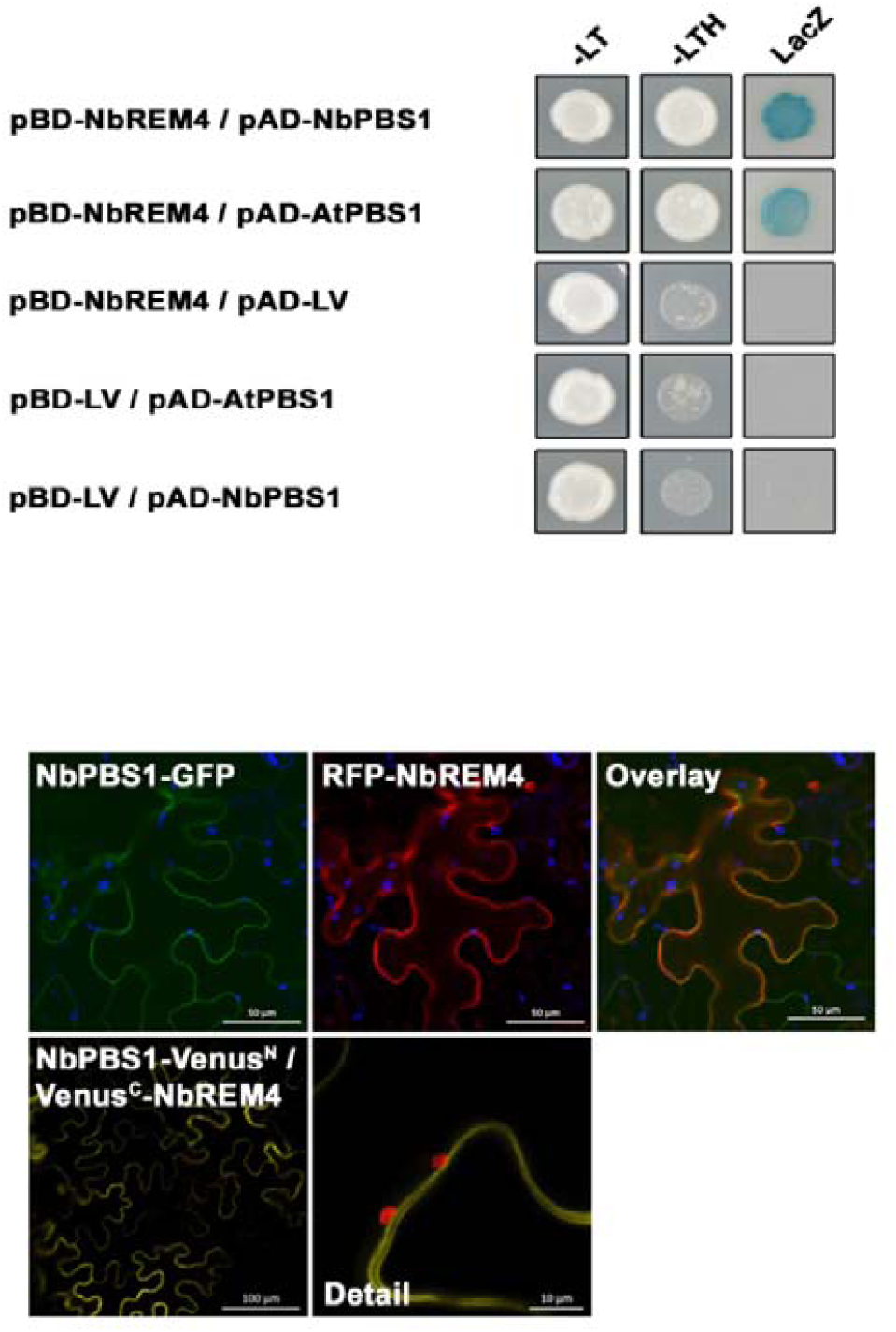
NbREM4 interacts with PBS1 in yeast two-hybrid assays and *in planta*. **A,** NbREM4 fused to the GAL4 DNA binding domain (BD) was expressed in combination with PBS1 fused to the GAL4 activation domain (AD) in yeast strain Y190. Cells were grown on selective media before a LacZ filter assay was performed. The empty BD vector or AD vector, respectively, served as negative control. NbPBS1, *N. benthamiana* PBS1; AtPBS1, *A. thaliana* PBS1. – LT, yeast growth on medium without Leu and Trp. – HLT, yeast growth on medium lacking His, Leu, and Trp, indicating expression of the *HIS3* reporter gene. LacZ, activity of the lacZ reporter gene. **B,** Co-localization of NbPBS1-GFP and RFP-NbREM4 in *N. benthamiana* leaf epidermal cells. NbPBS1-GFP was co-expressed with RFP-NbREM4 by Agrobacterium-infiltration. The green fluorescence (GFP), red fluorescence (mCherry) and chlorophyll autofluorescence (blue) were monitored separately to prevent cross-talk of the fluorescence channels and the resulting fluorescence images were merged. **C,** BiFC *in planta* interaction of NbPBS1 with NbREM4. YFP confocal microscopy images show *N. benthamiana* leaf epidermal cells transiently expressing NbPBS1-Venus^N^ in combination with Venus^C^-NbREM4. A close-up of the same cells shows that the YFP fluorescence produced by the interaction of NbPBS1-Venus^N^ with Venus^C^-NbREM4 aligns with the plasma membrane.

To assess the extent of co-localization of NbREM4 and PBS1 inside plant cells, NbREM4 N-terminally tagged with the red fluorescence protein (RFP) and NbPBS1-GFP were co-expressed in leaves of *N. benthamiana* using *Agrobacterium*-infiltration. Confocal imaging 2 dpi revealed that green as well as red fluorescence aligned with the region representing the PM and both fluorescence signals displayed substantial overlap indicative for co-localization of both proteins (Figure 7B). We applied the BiFC approach to analyse whether the observed Y2H interactions occur in living plant tissues/cells. Transient co-expression of NbPBS1-Venus^N^ together with Venus^C^-NbREM4 induced a strong fluorescence signal along the PM not engulfing the chloroplasts (Figure 7C). Control experiments indicate that the BiFC signal is specific (Supplementary Figure S5). In summary, these experiments suggest that NbREM4 specifically interacts with PBS1 at the PM of plant cells.

### PBS1 phosphorylates NbREM4 in vitro

Arabidopsis PBS1 belongs to subgroup VII of RLCK of which several members have been shown to play redundant roles in mediating PTI responses upon phosphorylation by upstream PRRs (Yamada et al., 2016; Bi et al., 2018; Rao et al., 2018). Based on protein mobility shifts on SDS-PAGE, PBS1 has been shown to undergo flg22 dependent phosphorylation in Arabidopsis protoplasts, suggesting an activation of the kinase during PTI signaling (Lu et al., 2010; Zhang et al., 2010). However, downstream phosphorylation targets of PBS1 have so far not been identified. Given that PBS1 interacts with NbREM4 and that remorins have been described as phosphoproteins (Reymond et al., 1996; Benschop et al., 2007; Marín and Ott, 2012; Toth et al., 2012), we explored the possibility of NbREM4 being a phosphorylation substrate of PBS1. To this end both proteins were recombinantly produced in *E. coli* and subjected to an *in vitro* phosphorylation assay using radiolabeled ATP. As indicated by their autophosphorylation activity both NbPBS1 and AtPBS1 recombinant proteins appeared to be active *in vitro*. Under our experimental conditions AtPBS1 reproducibly produced a stronger autophosphorylation signal than NbPBS1, indicating higher activity of the recombinant protein from Arabidopsis. Thus, we used AtPBS1 for the *in vitro* phosphorylation of NbREM4. Only when MBP-AtPBS1 and MBP-NbREM4 were present in the same assay mix an additional prominent signal appeared that corresponds to the MBP-NbREM4 protein band, indicating phosphorylation of NbREM4 by AtPBS1 *in vitro* (Figure 8A). The addition of MBP alone produced no additional signal, demonstrating phosphorylation of the NbREM4 portion within the MBP-NbREM4 fusion protein.

**Figure 8.**
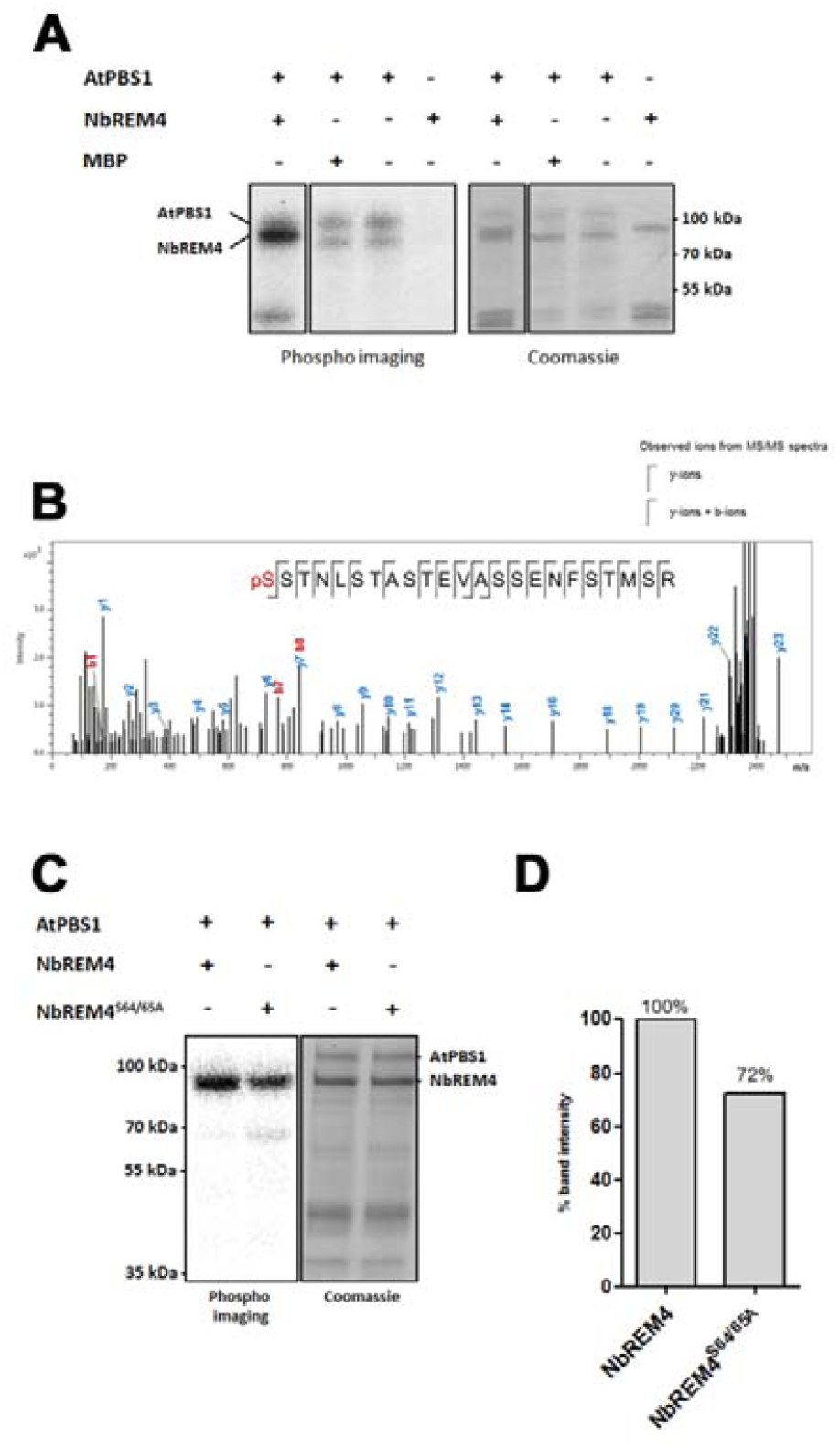
PBS1 phosphorylates NbREM4 *in vitro*. **A,** *In vitro* phosphorylation assay of NbREM4 by AtPBS1. Recombinant proteins were incubated in the presence of ^32^P-ATP and the mixture was subsequently resolved by SDS-PAGE. MBP (maltose-binding protein) served as a negative control. Left, autoradiogram; right, coomassie blue stain for protein visualization. **B,** MALDI-TOF MS/MS spectrum of an ion at *m/z* 2474.03 derived from *in vitro* phosphorylated NbREM4. Peptide fragmentation provides strong evidence for the sequence ^64^S to ^86^R of the NbREM4 polypeptide with phosphorylation occurring at ^64^S (inset). Intact y-ions are labeled in blue, intact b-ions are labeled in red. C, In vitro phosphorylation of NbREM4^S64/64A^ compared to the wild type NbREM4 protein. Recombinant proteins were incubated in the presence of ^32^P-ATP and the mixture was subsequently resolved by SDS-PAGE. Left, autoradiogram; right, coomassie blue stain for protein visualization. **D,** Densitometric analysis of NbREM4^S64/64A^ phosphorylation using ImageJ.

In order to map NbREM4 phosphorylation sites, phosphorylation reactions were repeated under non-radioactive conditions and protein from gel excised MBP-NbREM4 bands was subjected to MALDI-TOF MS/MS analysis. This identified a singly phosphorylated peptide comprising the amino acids 64 – 87 of the NbRemorin4 protein (relative to the start M of the native NbREM4 protein, Figure 8B). The peptide contained several serine and threonine residues that potentially could serve as phosphoacceptor sites. Fragmentation analysis suggested phosphorylation at one of the N-terminal residues of the phosphopeptide located within the double serine motif S64 (62 % probability) or S65 (35 % probability). We created an MBP-NbREM4^S64/65A^ variant that carries a serine to alanine substitution at the potential phosphorylation site and used the purified recombinant protein in an *in vitro* kinase assay with AtPBS1. This still yielded a signal corresponding to MBP-NbREM4^S64/65A^, indicating that the serine to alanine substitution at this position did not abolish phosphorylation (Figure 8C). However, a densitometric analysis of signal intensity revealed a reduction of phosphorylation by app. 30 % (Figure 8D). This suggests that phosphorylation of NbREM4 by AtPBS1 at either S64 or S65 occurs but that the protein contains additional phosphorylation sites that escaped the MALDI-TOF MS/MS analysis. The presence of additional phosphosites in NbREM4 is in accordance with an *in silico* phosphosite prediction using NetPhos3.1 (http://www.cbs.dtu.dk/services/NetPhos/) which predicts at least 28 putative serine or threonine phosphorylation sites with high confidence (Supplementary Figure S8).

Taken together, these data suggest that PBS1 phosphorylates NbREM4 at multiple sites *in vitro* and thus opens the possibility that NbREM4 is also a phosphorylation target of PBS1 *in planta*.

### NbREM4 is not acetylated by HopZ1a in vitro

HopZ1a possess acetyltransferase activity and acetylation of target proteins has been associated with virulence and avirulence functions of the effector (Lee et al., 2012b; Jiang et al., 2013; Lewis et al., 2013). In order to investigate whether the HopZ1a interaction partner NbREM4 also serves as an acetylation target an *in vitro* acetylation assay was conducted using *E. coli* produced proteins. Purified recombinant proteins were incubated with ^14^C-acetyl-coenzyme A and 100 nM inositol-hexakisphosphate (IP6) for 1 h at 30°C. Subsequently the proteins were separated by SDS-PAGE analysis followed by autoradiography. IP6 is a eukaryotic cofactor that stimulates the acetyltransferase activity of effectors in the YopJ family, including HopZ1a (Lee et al., 2012b).

Auto-acetylation of GST-HopZ1a was readily detected, indicating enzymatic activity of the recombinant protein (Supplementary Figure S9). However, no clear signs of an acetylation of the MBP-NbREM4 band could be detected suggesting that the remorin does not constitute an acetylation target of HopZ1a *in vitro*.

### NbREM4 relocates to PM microdomains after flg22 treatment

Previous reports have localized individual remorin isoforms from different plant species to PM membrane microdomains either in a constitutive or stimulus dependent manner (Raffaele et al., 2009; Lefebvre et al., 2010; Demir et al., 2013; Jarsch et al., 2014). Transient expression of GFP-NbREM4 in leaves of *N. benthamiana* did not reveal clear signs of localization to distinct membrane subdomains (Figure 9). Given the fact that NbREM4 could potentially be involved in PAMP signaling at the PM we investigated whether flg22 treatment would have any influence on GFP-NbREM4 PM localization. Indeed, we could observe that one hour after treatment with flg22 GFP-NbREM4 fluorescence was increasingly found in distinct and immobile membrane microdomains that were distributed over the entire inner PM leaflet of treated cells (Figure 9).

**Figure 9.**
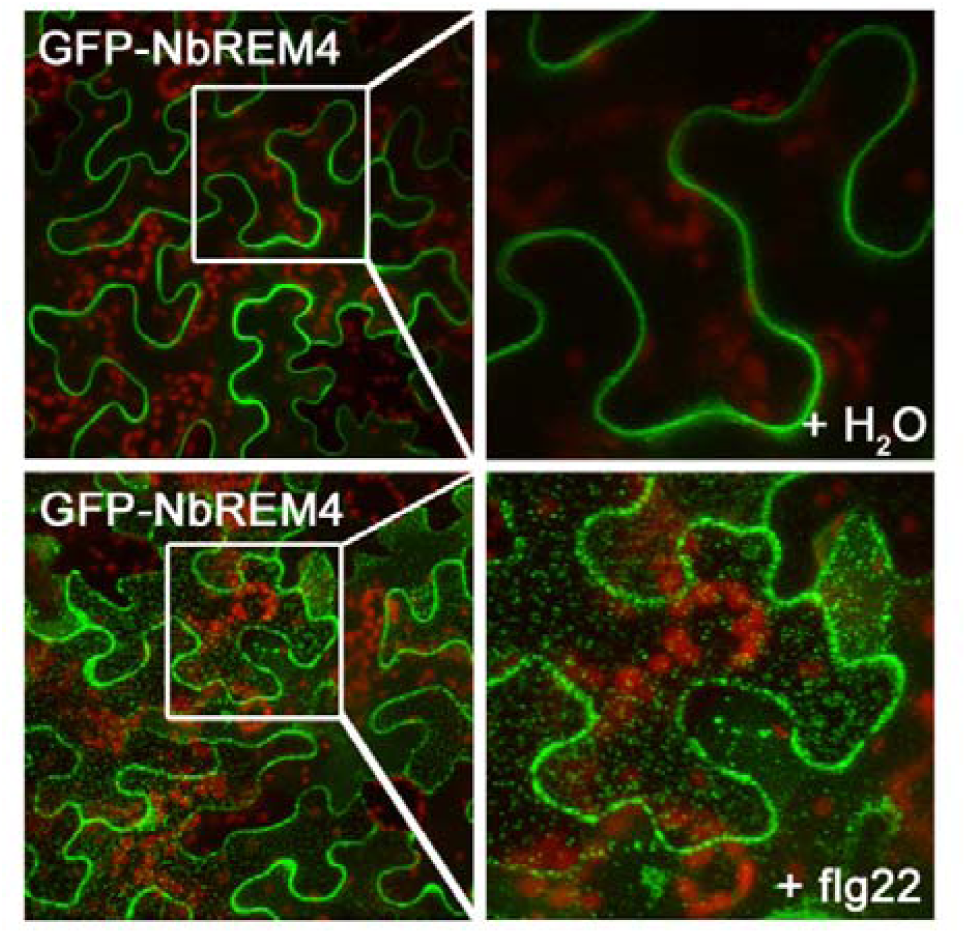
Relocation of NbREM4 upon flg22 treatment into punctuate structures. Confocal micrographs of *N. benthamiana* leaf epidermal cells transiently expressing GFP-NbREM4 challenged flg22 as compared to a H_2_O control. The flg22 treatment was performed 48 hpi and 60 min before imaging. Left, overview; right, magnification of the inset indicated on the left.

### Possible role of NbREM4 in HopZ1a dependent ETI

HopZ1a has been shown to elicit an HR in *N. bemthamiana*, indicative for its recognition through an R protein (Ma et al., 2006). However, Baudin et al. (2017) recently found that inducible expression of HopZ1a in *N. benthamiana* did not induce an HR. In Arabidopsis, HopZ1a recognition requires the canonical CC-type NLR protein ZAR1 (HOPZ-ACTIVATED RESISTANCE1) (Lewis et al., 2010). Detection of HopZ1a by AtZAR1 requires AtZED1 (HOPZ-ETI-DEFICIENT1), a receptor-like cytoplasmic kinase (RLCK) belonging to the RLCK clade XII-2 family that acts as decoy guarded by AtZAR1 to senses the activity of HopZ1a (Lewis et al., 2013). Given the discrepancy in the observed HopZ1a overexpression phenotypes, we initiated a set of experiments to further investigate a HopZ1a dependent HR in *N. benthamiana* and explore the possible involvement of NbREM4 and PBS1 in this process. First, we assessed whether under our experimental conditions expression of HopZ1a leads to HR development in *N. benthamiana*. To this end, HopZ1a, HopZ1a^C/A^ and the *Xanthomonas campestris* pv. *vesicatoria* T3E AvrRxv were transiently expressed in *N. benthamiana* leaves alongside with an empty vector (EV) control. As compared to the EV control and the catalytically inactive HopZ1a^C/A^ variant, HopZ1a infiltrated areas showed clear signs of cell death at 2 dpi that were even more prominent than those triggered by AvrRxv (Figure 10A). In accordance with the observed leaf phenotype, ion leakage from HopZ1a infiltrated tissue was significantly increased at 2 dpi while expression of HopZ1^C/A^ did not increase electrolyte efflux (Figure 10B). A western blot analysis confirmed expression of all proteins tested (Figure 10C). Thus, transient expression using *Agrobacterium*-infiltration triggers HR-like symptoms and membrane damage in *N. benthamiana*. In order to investigate whether the observed HopZ1a-dependent phenotype would require the R protein ZAR1, we cloned a fragment of the *N. benthamiana ZAR1* orthologue (Baudin et al., 2017) into a vector for virus-induced gene silencing (VIGS). Strong down-regulation of *NbZAR1* could be confirmed in *N. benthamiana* plants 2 weeks after infiltration with the VIGS vectors as compared to the *GFP*-VIGS (*GFP*-VIGS) control (Figure 11A). Transient expression of HopZ1a in *GFP*-VIGS plants led to the previously observed HR symptoms at 2 dpi while HopZ1aC/A did not cause phenotypic changes (Figure 11B). In contrast, HopZ1a expression in *NbZAR1*-VIGS plants did not yield visible signs of HR at 2 dpi (Figure 11B). Protein expression in all tissues analyzed was confirmed by western blotting (Figure11C). Ion leakage measurement revealed that HopZ1a did not cause a significant increase in membrane damage in *NbZAR1*-VIGS plants as opposed to the *GFP*-VIGS control (Figure 11D). Taken together, the data suggest that similar to the situation in Arabidopsis (Lewis et al., 2010), the HopZ1a-dependent HR in *N. benthamiana* depends on its recognition by NbZAR1.

**Figure 10.**
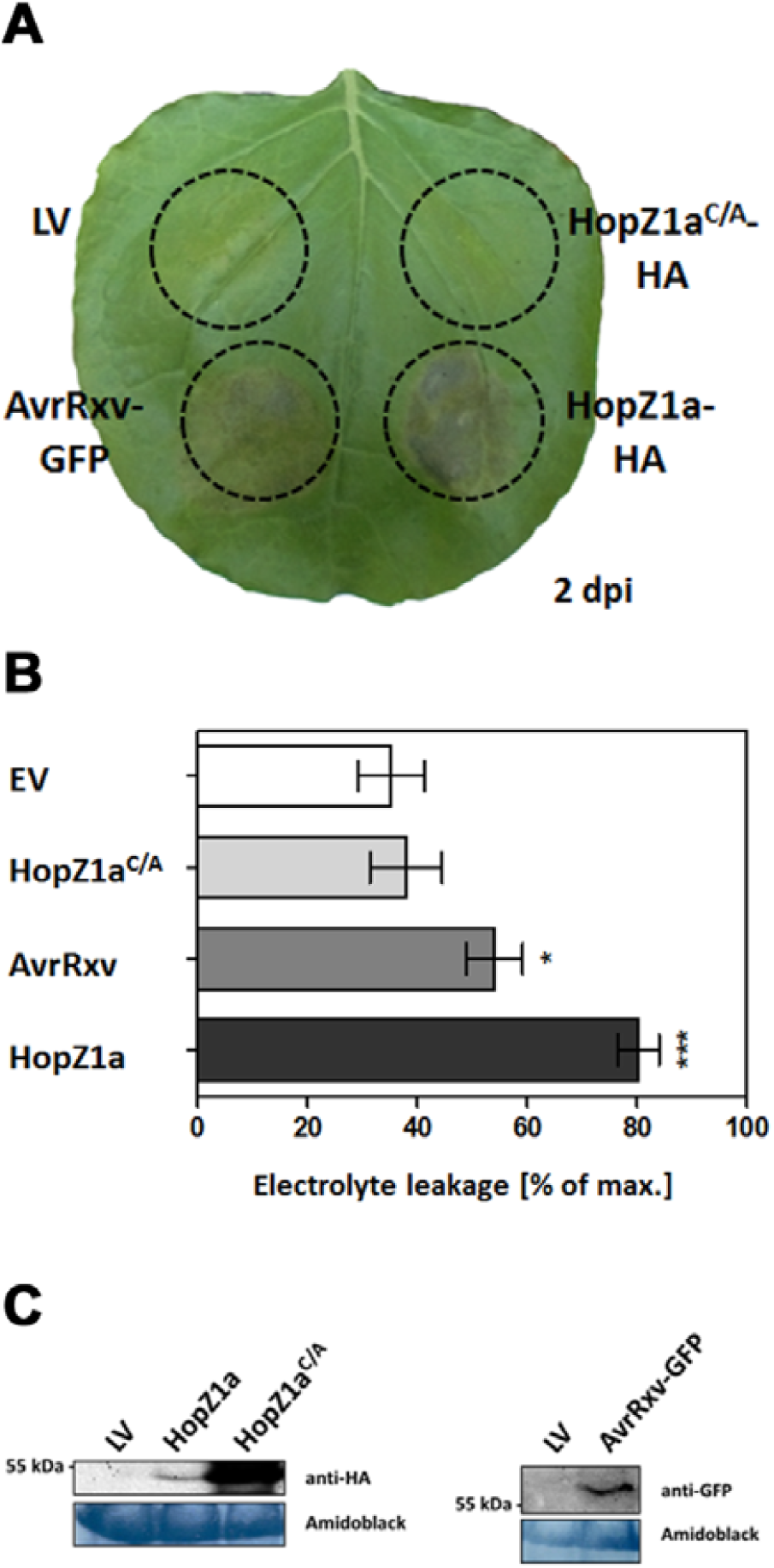
Transient HopZ1a expression induces an HR in *N. benthamiana* leaves. **A,** *Agrobacterium*-mediated transient expression assay in *N. benthamiana*. The constructs indicated were inoculated into the leaf areas delineated by the dashed line. The picture was taken 48 hpi. **B,** Ion leakage was measured in plants transiently expressing the constructs indicated 48 hpi with *Agrobacteria*. Bars represent the average ion leakage measured for triplicates of six leaf disks each, and the error bars indicate SD. The asterisk indicates a significant difference (\*\**P* < 0.05,\*\*\**P* < 0.01) based on results of a Student’s *t*-test. **C,** Protein immunoblot of HopZ1a-HA, HopZ1^C/A^-HA and AvrRxv-GFP, verifying protein expression. Samples were collected 48 hpi and probed with α-HA antibody or α-GFP antibody, respectively. Amido-black staining is included to show equal loading of the samples.

**Figure 11.**
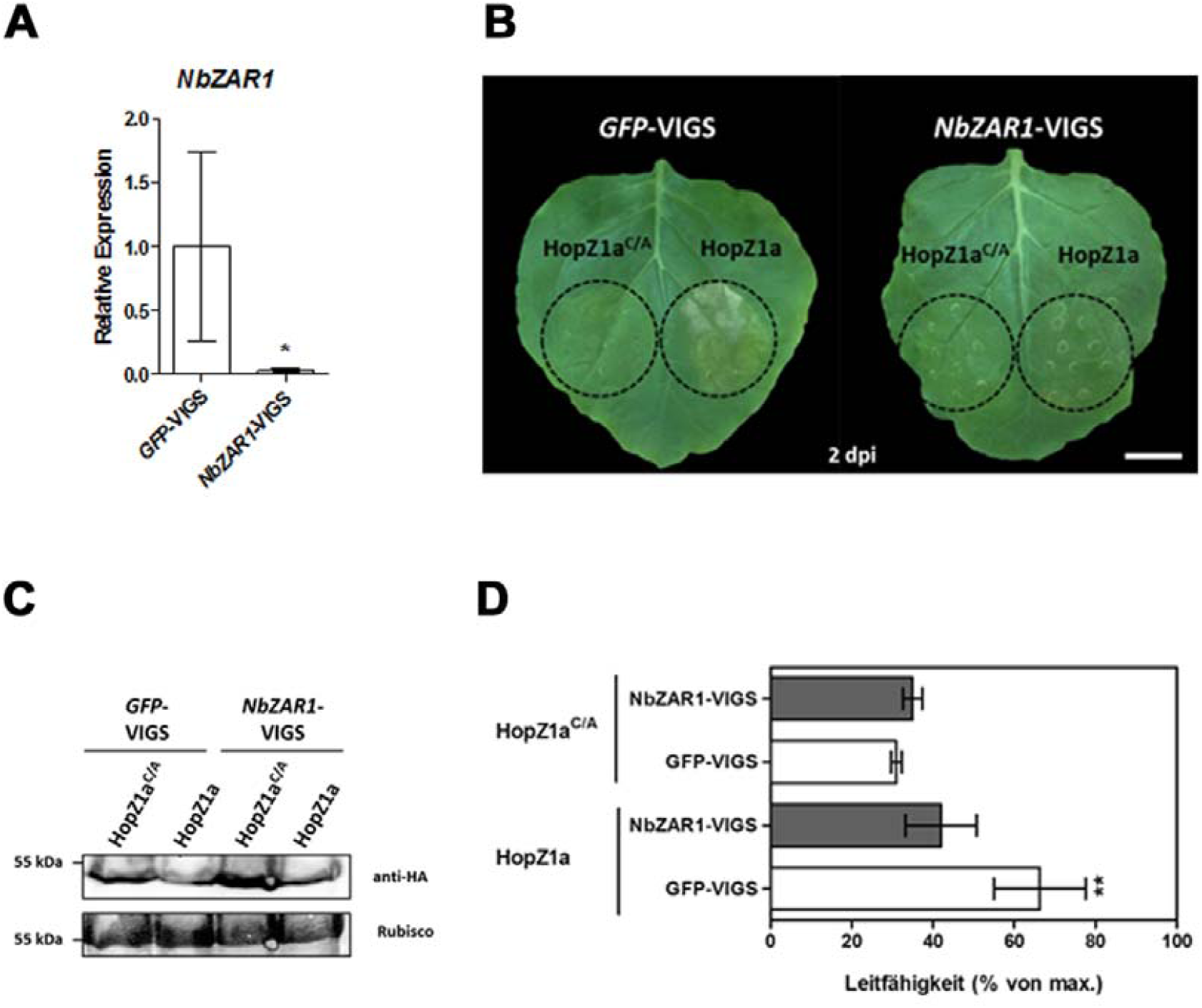
Down-regulation of ZAR1 expression abolishes HopZ1a triggered HR. **A,** quantitative real-time RT-PCR analysis of *ZAR1* down-regulation in *NbZAR1*-VIGS *N. benthamiana* plants. Total RNA was isolated from leaves of VIGS plants 14 dpi and subjected to cDNA synthesis followed by qRT-PCR. Bars represent the mean of at least three biological replicates ± SD. The asterisk indicates a significant difference (*P < 0.05) based on results of a Student’s t-test. **B,** Leaves of *NbZAR1*-VIGS and *GFP*-VIGS plants were infiltrated with the constructs indicated by the dashed line. **C,** Protein immunoblot of HopZ1a-HA and HopZ1^C/A^-HA, verifying protein expression VIGS plants. Samples were collected 2 dpi and probed with an α-HA antibody. Amido-black staining is included to show equal protein loading. **D,** Ion leakage was measured in VIGS plants transiently expressing the constructs indicated on the left 48 hpi. Bars represent the average ion leakage measured for triplicates of six leaf discs each, and the error bars indicate SD. The asterisk indicates a significant difference (\*\**P* < 0.01) based on results of a Student’s *t*-test.

We next sought to explore whether the HopZ1a interaction partner NbREM4 contributes to HopZ1a-asssociated HR in *N. benthamiana*. Therefore, we used the VIGS system to repress NbREM4 expression and subsequently expressed HopZ1a in NbREM4-silenced plants. NbREM4 expression was drastically reduced in NbREM4-VIGS plants, indicating efficient silencing (Supplementary Figure S10A). However, *NbREM4*-VIGS plants displayed similar HR symptoms and electrolyte leakage upon HopZ1a expression as the *GFP*-VIGS control (Supplementary Figure S10), suggesting that down-regulation of NbREM4 does not affect the ability of HopZ1a to elicit an HR.

## Discussion

During infection, bacterial T3Es translocated by the type III secretion system play a central role in the manipulation of the host cellular machinery in favor of the pathogen. In general, these T3Es are essential for pathogen virulence by interfering with plant processes involved in defense responses (Jones and Dangl, 2006). In order to interact with the correct host target and exert their function, T3Es often show a specific localization within the host cell (Hicks and Galán, 2013). A recent survey suggests that in case of *P. syringae* more than half of the T3Es characterized to date target host proteins that localize to the host cell’s plasma membrane (Khan et al., 2018a). This highlights the importance of the PM as the interface of pathogen recognition and initiation of immunity (Hoefle and Hückelhoven, 2008). Accordingly, a number of effector proteins carry lipid modifications such as acylation (also called palmitoylation), myristoylation, and prenylation that enable host membrane association after translocation into the cytosol (Hicks and Galán, 2013). The *P. syringae* T3E HopZ1a is an acetyltransferase that requires N-terminal myristoylation for PM targeting and this modification is required for its virulence and avirulence function in Arabidopsis and soybean (Lewis et al., 2008). Several distinct molecular targets of HopZ1a have been identified not all of which are known to locate to the PM (Zhou et al., 2011; Lee et al., 2012b; Jiang et al., 2013). However, HopZ1a has been shown to interfere with PM and cell wall-associated defense such as callose deposition, the ROS burst and MAP kinase activation (Lee et al., 2012b; Lewis et al., 2014). This is partially explained by its ability to cause microtubule destruction through acetylation of tubulin (Lee et al., 2012b).

Here, we identified the remorin protein NbREM4 from *N. benthamiana* as a novel interaction partner of HopZ1a in plants. The obtained evidence suggests that the HopZ1a NbREM4 interaction occurs at the PM and does not require additional proteins. Remorins are plant specific proteins with multiple roles in plant microbe interactions, including those with viruses (Raffaele et al., 2009; Perraki et al., 2014; Son et al., 2014; Fu et al., 2018), oomycetes (Bozkurt et al., 2014), and bacteria (Lefebvre et al., 2010; Toth et al., 2012; Liang et al., 2018). Interestingly, the Arabidopsis remorins AtREM1.2 and AtREM1.3 have been identified to interact with the Pseudomonas T3E HopF2 in vivo by two independent methods (Hurley et al., 2014; Khan et al., 2018b). Although the functional significance of this interaction is currently unknown it might point to broader role of remorins in T3E function in plants. All described remorins associate with the PM in dynamic microdomains known as membrane rafts (Jarsch and Ott, 2011; Jarsch et al., 2014; Gronnier et al., 2017). In mammals and plants, these rafts have been suggested to constitute assembly platforms for signal transduction, pathogen infection, and other processes (Simons and Gerl, 2010; Simon-Plas et al., 2011). It has been shown that *Medicago truncatula* nodulation-induced remorin MtSYMREM1 interacts with at least three RLKs NFP, LYK3, and DMI that are essential for root nodule symbiosis (Lefebvre et al., 2010). Infection dependent induction of MtSYMREM1 results in recruitment of ligand activated LYK3 and its stabilization within membrane subdomains to prevent endocytosis of the receptor, a function required for successful rhizobial infection (Liang et al., 2018). We observed relocalization of GFP-NbREM4 to distinct PM subdomains resembling lipid rafts after treatment with flg22. A similar redistribution into specific PM subdomains upon elicitation with flg22 has also been observed for the flg22 receptor FLS2 (Keinath et al., 2010). This supports a model in which PAMP-induced signaling may require defined membrane domains that provide physical rafts for receptor-scaffold and other protein-protein interactions possibly involving NbREM4. Phosphorylation could be one mechanism which regulates membrane sublocalization and/or protein-protein interactions of NbREM4. Many remorins have been shown to be phosphorylated in a constitutive or stimulus dependent manner (Jarsch and Ott, 2011). In a large scale proteomic approach to identify differentially phosphorylated proteins involved in early PAMP, AtREM1.3 was shown to be phosphorylated in an flg-22 dependent manner while other remorin isoforms displayed constitutive phosphorylation (Benschop et al., 2007). The potato remorin StREM1.3 is differentially phosphorylated by a PM associated protein kinase upon the perception of polygalacturonic acid (Reymond et al., 1996; Jarsch and Ott, 2011). Generally, phosphorylation of remorins appears to be restricted to serine and threonine residues that are exclusively located within the lowly conserved N-terminal region of the protein (Marín and Ott, 2012). The N-terminal domain of remorins is intrinsically disordered and that phosphorylation within this region has the capacity to alter the interaction of remorins with other proteins (Marín et al., 2012). We could show that NbREM4 is phosphorylated on several residues, including serine 64 within the unconserved N-terminal domain, by the PM localized RLCK PBS1 *in vitro*. PBS1 belongs to subfamily VII of RLCKs whose members have been implicated in several aspects of PTI, including activation of MAP signaling, generation of ROS, and transcriptional reprogramming (Lin et al., 2013; Bi et al., 2018; Lal et al., 2018; Rao et al., 2018). The role of PBS1 in PTI signaling is not entirely clear. It has been shown that flg22 treatment induces phosphorylation of PBS1 in Arabidopsis as well as in wheat (Lu et al., 2010; Sun et al., 2017). However, genetic studies indicate that at least in Arabidopsis PBS1 only plays a minor role PTI signaling as compared to other members of the RLCK VII family (Zhang et al., 2010). Whether PBS1 in other plants is more important to trigger PAMP induced defense responses is currently unknown. NbREM4 does not seem to interact with other tested members of the RLCK VII clade in yeast than PBS1. However, this does not exclude phosphorylation of NbREM4 by other kinases *in vivo*. Phosphorylation of NbREM4 by an upstream RLCK, like PBS1, could then lead to activation of downstream defense responses for instance by altering the ability of NbREM4 to interact with other proteins and thereby affecting their activity. Along that line, we could observe that transient overexpression of NbREM4 in leaves of *N. benthamiana* led to the development of leaf chlorosis associated with membrane damage and the induction of defense related genes. Thus, overexpression of NbREM4 might dominantly activate downstream defense responses. However, we currently cannot exclude a general cytotoxic effect of NbREM4 through interference with other vital cellular processes. The unequivocal demonstration that NbREM4 phosphorylation is required for relocalization of the protein into membrane microdomains or that it alters protein-protein interactions of NbREM4 downstream immune related targets requires further experimentation.

The virulence function of HopZ1a has been associated with its acetyltransferase activity and acetylation of host target proteins by HopZ1a is thought to interfere with their function in immunity related processes (Zhou et al., 2011; Lee et al., 2012a; Jiang et al., 2013). The results of *in vitro* acetylation assays do not suggest acetylation of NbREM4 by HopZ1a and thus it is currently unknown whether NbREM4 constitutes a genuine effector target for HopZ1a or whether it might have some auxiliary function. Remorins have thus far not been described as target proteins for bacterial T3Es but given their extensive involvement in plant-microbial interactions interference with remorin function could potentially have a wide impact on PTI related defense outputs. Alternatively, HopZ1a could use NbREM4 as an auxiliary factor that facilitates either the indirect interaction with other proteins or the localization to specific membrane subdomains related to HopZ1a function. RLCKs like PBS1 could constitute these indirect targets and future experiments have to investigate whether PBS1 or other NbREM4 interacting proteins are substrates for HopZ1a acetylation and hence whether interaction of HopZ1a with NbREM4 is related to the virulence function of the effector.

In Arabidopsis, HopZ1a is recognized by the NLR ZAR1 to trigger ETI. Recognition depends on the pseudokinase ZED1 which acts as a decoy for HopZ1a mediated acetylation to activate ZAR1 (Lewis et al., 2013). Recent evidence suggests that the ZAR1 immune pathway is largely conserved in *N. benthamiana* (Baudin et al., 2017). Our observation that transient expression of HopZ1a in *N. benthamiana* triggers an ETI response is in line with previous reports (Ma et al., 2006) but in contrast to a recent study by Baudin et al. (2017) who observed a HopZ1a dependent HR in *N. benthamiana* only upon coexpression of the effector with ZED1 from Arabidopsis. The discrepancy between these two observations is likely due to differences in the experimental setup. While in Ma et al. (2006) and the study described here a strong constitutive CaMV35S promoter was used to drive HopZ1a expression in *N. benthamiana* leaves, Baudin et al. (2017) used an inducible system that might lead to differences in expression kinetics and protein amounts of the effector and thus could affect the outcome of the overexpression. As previously observed (Baudin et al., 2017), silencing of ZAR1 abolished the HopZ1a triggered HR in *N. benthamiana* confirming effector recognition by this NLR also in this species. Based on the current model of HopZ1a recognition by ZAR1 in Arabidopsis, this implies that HopZ1a likely interacts and modifies a protein that is guarded by ZAR1 in *N. benthamiana*. A ZED1 orthologue in *N. benthamiana* has so far not been characterized and thus potential guardees of NbZAR1 are currently unknown. Recent data suggest that ZAR1 in Arabidopsis interacts with multiple guardees beyond ZED1, including ZRK1 (ZED-related 1) leading to the recognition of AvrAC from *Xanthomonas campestris* pv. *campestris* (Wang et al., 2015) and ZRK3 required for the perception of HopF2a from *Pseudomonas syringae* (Seto et al., 2017). Thus, it is possible that in *N. benthamiana* ZAR1 guards various members of the RLCK family. In Arabidopsis, PBS1 is guarded by the NLR RPS5 which activates HR upon proteolytic cleavage of PBS1 by the *P. syringae* T3E AvrPphB (Shao et al., 2003), thus providing precedent for a role of PBS1 in ETI. Recognition of AvrAC in Arabidopsis requires PBS-like 2 (PBL2) in addition to ZRK1 and the current model suggests that ZAR1 forms a stable complex with ZRK1, which specifically recruits PBL2 when the latter is uridylylated by AvrAC to subsequently trigger ZAR1-mediated immunity (Wang et al., 2015). Thus, modification of PBL2 by AvrAC is indirectly recognized by the ZAR1/ZRK1 immune complex. Silencing of NbREM4 did not affect HopZ1a triggered HR in *N. benthamiana*, arguing against a direct or indirect involvement of this to protein in HopZ1a recognition.

In summary, the present study shows that HopZ1a directly interacts with NbREM4 in *N. benthamiana* and transient overexpression of NbREM4 point towards a role of this remorin in induced defense responses. This notion is corroborated by the finding that NbREM4 interacts and is phosphorylated by the immune related RLCK PBS1. Future studies have to clarify whether the interaction with HopZ1a interferes with this possible role of NbREM4 in defense. Virus-induced gene silencing studies indicate that neither NbREM4 nor PBS1 play a role in HopZ1a triggered ETI in *N. benthamiana* favoring a role of these proteins as virulence targets for this T3E. The finding that HopZ1a triggers a ZAR1-dependent HR in *N. benthamiana* in absence of the Arabidopsis decoy protein ZED1 further extends on the observation concerning conservation of ZAR1-mediated HopZ1a recognition in this species (Baudin et al., 2017).

## Material and Methods

### Plant material and growth conditions

Tobacco plants (*Nicotiana benthamiana*) were grown in soil in a climate chamber with daily watering, and subjected to a 16 h light/8 h dark cycle (25°C: 21°C) at 300 µmol m^−2^ s^−1^ light and 75% relative humidity.

### Yeast Two-Hybrid Analysis

Yeast two-hybrid techniques were performed according to the yeast protocols handbook and the Matchmaker GAL4 Two-hybrid System 3 manual (both Clontech, Heidelberg, Germany) using the yeast reporter strains AH109 and Y187. The entire HopZ1a coding region was amplified by PCR using the primers listed in Table S1 and inserted in the pGBT-9 vector generating a fusion between the GAL4 DNA-binding domain (BD). The yeast strain Y187 carrying the BD-HopZ1a construct was mated with AH109 cells pre-transformed with a library derived from tobacco (*Nicotiana tabacum*) source leaves (Börnke, 2005). Diploid cells were selected on medium lacking Leu, Trp, and His supplemented with 4 mM 3-aminotriazole. Cells growing on selective medium were further tested for activity of the *lacZ* reportergene using filter lift assays. Library plasmids from *his3/lacZ* positive clones were isolated from yeast cells and transformed into *E. coli* before sequencing of the cDNA inserts. Direct interaction of two proteins was investigated by cotransformation of the respective plasmids in the yeast strain AH109, followed by selection of transformants on medium lacking Leu and Trp at 30°C for 3 days and subsequent transfer to medium lacking Leu, Trp and His for growth selection and *lacZ* activity testing of interacting clones.

### Plasmid construction

To generate plasmids containing the corresponding gene of interest, the entire open reading frame was amplified by PCR from *Arabidopsis* cDNA using the primers listed in Supplementary Table SX. The resulting fragments were inserted into the pENTR-D/TOPO vector according to the manufacturer’s instructions (Thermo) and verified by sequencing. For yeast two-hybrid analysis, fragments were recombined into Gateway®-compatible versions of the GAL4-DNA binding domain vector pGBT-9 and the activation domain vector pGAD424 (Clontech) using L/R-clonase (Thermo). To generate translational fusions between NbREM4 and the green fluorescent protein (GFP) coding sequences were inserted into the vector pK7FWG2 (Karimi et al., 2002). Constructs for bi-molecular complementation analysis are based on Gateway®-cloning compatible versions of pRB-C-Venus^N173^and pRB-C-Venus^C155^.

### Agrobacterium*-infiltration*

For infiltration of *N. benthamiana* leaves, *A. tumefaciens* C58C1 was infiltrated into the abaxial air space of 4-to 6-week-old plants, using a needleless 2-ml syringe. Agrobacteria were cultivated overnight at 28°C in the presence of appropriate antibiotics. The cultures were harvested by centrifugation, and the pellet was resuspended in sterile water to a final optical density at (OD_600_) of 1.0. The cells were used for the infiltration directly after resuspension. Infiltrated plants were further cultivated in the greenhouse daily watering, and subjected to a 16 h light/8 h dark cycle (25°C/21°C) at 300 µmol m^−2^ s^−1^ light and 75% relative humidity.

### In vitro *pull-down*

Recombinant MBP-NbREM4 from *Escherichia coli* (BL21 DE3, New England Biolabs) lysates was immobilized on amylose resins (New England Biolabs) and incubated for 1 h at 4°C with total protein lysates from cell expressing GST-HopZ1a or GST-NbPBS1. Proteins were eluted, and analyzed by immunoblotting using either anti-GST antibody (Sigma) or anti-MBP antibody (NEB).

### Western blotting

Leaf material was homogenized in sodium-dodecyl sulphate-polyacrylamide gel electrophoresis (SDS-PAGE) loading buffer (100 mM Tris-HCl,pH 6.8;9% β-mercaptoethanol, 40% glycerol, 0.2% bromophenol blue, 4% SDS) and, after heating for 10 min at 95°C, subjected to gel electrophoresis. Separated proteins were transferred onto nitrocellulose membrane (Porablot, Machery und Nagel, Düren, Germany). Proteins were detected by an anti-HA-peroxidase high-affinity antibody (Roche) or anti-GFP antibody (Roche) via chemiluminescence (GE Healthcare) using a myECL imager (Thermo).

### Bimolecular fluorescence complementation (BiFC)

Constructs were transformed into *A. tumefaciens* C58C1 and transiently expressed by Agrobacterium-infiltration in *N. benthamiana*. The BiFC-induced YFP fluorescence was detected by CLSM (LSM510; Zeiss) 48 hpi. The specimens were examined using the LD LCI Plan-Apochromat 253/0.8 water-immersion objective for detailed images with excitation using the argon laser (458-or 488-nm line for BiFC and chlorophyll autofluorescence). The emitted light passed the primary beamsplitting mirrors at 458/514 nm and was separated by a secondary beam splitter at 515 nm. Fluorescence was detected with filter sets as follows: on channel 3, 530–560 band pass; and on channel 1, for red autofluorescence of chlorophyll.

### RNA Extraction and quantitative real-time PCR

Total RNA was isolated from leaf material and then treated with RNase-free DNase to degrade any remaining DNA. First-strand cDNA synthesis was performed from 2 µg of total RNA using Revert-Aid reverse transcriptase (Thermo). For quantitative RT-PCR, the cDNAs were amplified using SensiFAST SYBR Lo-ROX Mix (Bioline) in the AriaMx Realtime PCR System (Agilent Technologies) as previously described (Arsova et al., 2010). At least three biological repeats and three technical repeats were used for each analysis. The transcript level was standardized based on cDNA amplification of *NbActin* as a reference. Statistical analysis was performed using Student’s t test. Primers are provided in Supplemental Table S1.

### Virus-induced gene silencing

Virus-induced gene silencing in *N. benthamiana* was essentially carried out as described previously (Liu et al., 2002a; Liu et al., 2002b). In brief, a fragment of *N. benthamiana* NbREM4 was amplified by PCR using the primers indicated in Table S1 and cloned into pTRV2-Gateway using the Gateway^TM^ recombination system (Invitrogen) as described in the section ‘Construction of expression plasmids’. The plasmids were transformed into *A. tumefaciens* C58C1. A lower leaf of a 4-week-old *N. benthamiana* plant was co-infiltrated with a mixture of agrobacteria carrying either pTRV1 or pTRV2 containing the target sequence or a GFP negative control fragment as described previously (Liu et al., 2002b). Silenced plants were analyzed 14 d post infiltration.

### Ion leakage measurements

For electrolyte leakage experiments, triplicates of 1.76 cm^2^ infected leaf material were taken at different time points as indicated. Leaf discs were placed on the bottom of a 15-ml tube. Eight milliliters of deionized water was added to each tube. After 4 h of incubation in a rotary shaker at RT, conductivity was determined with a conductometer. To measure the maximum conductivity of the entire sample, conductivity was determined after boiling the samples for 1 h (Stall et al., 1974).

### In vitro Kinase assay

Purified MBP-NbREM4 or MBP-NbPBS1 (2 µg) were incubated in 20 µl of reaction buffer (10 mM HEPES pH 7.4, 2 mM MgCl_2_, 2 mM MnCl_2_, 0,2 mM DTT) containing 2 µCi of [γ-^32^P] ATP (Hartmann Analytics). Reactions were incubated at 30°C for 1 h. The reaction was stopped by adding 4 × Laemmli buffer and separate on a 4-12% polyacrylamide gel. The gel was stained with Coomassie and incorporated radiolabel was visualized by autoradiography.

### In vitro acetylation assay

Purified proteins (2 µg) were incubated in 20 µl acetylation buffer [100 nM inositol hexakisphosphate (IP6), 50 mM HEPES at pH 8.0, 10% (vol/vol) glycerol, and 5 mM DTT], supplemented with 0,1 µCi of [^14^C] acetyl-CoA (40-60 mCi/mmol; Hartmann Analytics) for 1 h at 30°C. The reactions were stopped by adding 4 × Laemmli buffer and separated on a 4-12% polyacrylamide gel. Gels were fixed in fixation solution [5% (vol/vol) methanol and 10% (vol/vol) acetic acid] for 30 min. Gels were then dried and placed in a phosphorimager cassette for 10 d at RT. The SDS/PAGE gel was run in duplicate and stained with Coomassie blue to visualize proteins.

### Mass Spectrometric analysis of phosphorylation sites

MALDI-TOF-MS/MS analyses were carried out on MBP-tagged NbREM4 in the presence of NbPBS1 and kinase buffer with non-radioactive ATP. The reaction was incubated for 1 h at RT, stopped by adding 4 × Laemmli buffer, and separated on a 4-12% polyacrylamide gel. Bands corresponding to the size of MBP-NbREM4 were excised from the gel and subjected to tryptic digestion as described earlier (Witzel et al., 2017). Phosphorylated peptides were enriched using the High-Select^TM^ Fe-NTA Phosphopeptide Enrichment Kit (ThermoFisher Scientific) following the manufacturer’s instructions. The resulting eluate was desalted using C_18_ ZipTip^®^ Pipette Tips (MerckMillipore). After mixing 1:1 with 2,5-dihydroxybenzoic acid (Bruker Daltonik GmbH, 20 mg mL^-1^ in 30:70 [v/v] acetonitrile:0.1% trifluoroacetic acid in water, supplemented with 1% H_3_PO_4_) as a matrix, the sample was spotted onto a ground steel target and allowed to dry at room temperature. Mass spectrometric experiments were performed using an ultrafleXtreme MALDI-TOF instrument (Bruker Daltonik) run in positive ionisation mode, controlled by flexControl v3.4 software (Bruker Daltonik). The acquired peptide mass fingerprint data was processed with flexAnalysis v3.4 (Bruker Daltonik) and matched to the amino acid sequence of MBP-tagged NbREM4 via MASCOT search engine (Matrix Science). Search parameters were: monoisotopic mass accuracy, one missed cleavage and allowed variable modification as phosphorylation (ST). Peptide tolerance for peptide mass fingerprinting (PMF) was 50 ppm and, for *de novo* sequencing, peptide tolerance was 50 ppm and 0.7 Da fragment tolerance. Acquisition of LIFT spectra was pursued for a peptide *m/z* 2474.03, where a putative phosphorylation was detected. Peptide mass fingerprinting spectra and corresponding LIFT spectra were calibrated using external calibration (Peptide Calibration Standard II, Bruker Daltonik).

## Supplemental data

The following supplemental materials are available:

**Supplemental Figure S1:** Alignment of remorin protein sequences.

**Supplemental Figure S2:** Phylogenetic tree of Arabidopsis remorins incorporating NbRemorin.

**Supplemental Figure S3:** Yeast two-hybrid assay to test for the interaction of HopZ1a with other remorin isoforms.

**Supplemental Figure S4:** NbREM4 forms homomers in yeast and *in planta*.

**Supplemental Figure S5:** Negative controls for HopZ1a/NbREM4 BiFC experiments.

**Supplemental Figure S6:** Alignment of PBS1 protein sequences.

**Supplemental Figure S7:** NbREM4/RLCK interactions in yeast.

**Supplemental Figure S8:** NetPhos analysis of the NbREM4 protein sequence to putative predict phosphorylation sites.

**Supplementary Figure S9:** *In vitro* acetylation assay of NbREM4 by HopZ1a.

**Supplementary Figure S10:** Virus-induced gene silencing of *NbREM4* does not affect HR induction by HopZ1a expression in *N. benthamiana.*

**Supplementary Table S1:** Oligonucleotides used in this study.

## Acknowledgements

This work was supported by grants from the Deutsche Forschungsgemeinschaft (BO1916/5-1 and BO1916/5-2) to F.B. The skillful technical help of Mandy Heinze, Susanne Jeserigk and Kerstin Bieler is gratefully acknowledged.

## References

Arsova, B., Hoja, U., Wimmelbacher, M., Greiner, E., Üstün, S., Melzer, M., Petersen, K., Lein, W., and Börnke, F. 2010. Plastidial thioredoxin z interacts with two fructokinase-like proteins in a thiol-dependent manner: evidence for an essential role in chloroplast development in Arabidopsis and Nicotiana benthamiana. Plant Cell 22:1498–1515.

Baudin, M., Hassan, J.A., Schreiber, K.J., and Lewis, J.D. 2017. Analysis of the ZAR1 Immune Complex Reveals Determinants for Immunity and Molecular Interactions. Plant Physiol 174:2038–2053.

Benschop, J.J., Mohammed, S., O’Flaherty, M., Heck, A.J., Slijper, M., and Menke, F.L. 2007. Quantitative phosphoproteomics of early elicitor signaling in Arabidopsis. Mol Cell Proteomics 6:1198–1214.

Bi, G., Zhou, Z., Wang, W., Li, L., Rao, S., Wu, Y., Zhang, X., Menke, F.L.H., Chen, S., and Zhou, J.-M. 2018. Receptor-Like Cytoplasmic Kinases Directly Link Diverse Pattern Recognition Receptors to the Activation of Mitogen-Activated Protein Kinase Cascades in Arabidopsis. The Plant Cell 30:1543–1561.

Börnke, F. 2005. The variable C-terminus of 14-3-3 proteins mediates isoform-specific interaction with sucrose-phosphate synthase in the yeast two-hybrid system. J Plant Physiol 162:161– 168.

Bozkurt, T.O., Richardson, A., Dagdas, Y.F., Mongrand, S., Kamoun, S., and Raffaele, S. 2014. The Plant Membrane-Associated REMORIN1.3 Accumulates in Discrete Perihaustorial Domains and Enhances Susceptibility to <em>Phytophthora infestans</em>. Plant Physiology 165:1005– 1018.

Büttner, D. 2016. Behind the lines-actions of bacterial type III effector proteins in plant cells. FEMS Microbiol Rev 40:894–937.

Cesari, S. 2018. Multiple strategies for pathogen perception by plant immune receptors. The New phytologist 219:17–24.

Demir, F., Horntrich, C., Blachutzik, J.O., Scherzer, S., Reinders, Y., Kierszniowska, S., Schulze, W.X., Harms, G.S., Hedrich, R., Geiger, D., and Kreuzer, I. 2013. Arabidopsis nanodomain-delimited ABA signaling pathway regulates the anion channel SLAH3. P Natl Acad Sci USA 110:8296– 8301.

Dodds, P.N., and Rathjen, J.P. 2010. Plant immunity: towards an integrated view of plant-pathogen interactions. Nat Rev Genet 11:539–548.

Fu, S., Xu, Y., Li, C., Li, Y., Wu, J., and Zhou, X. 2018. Rice Stripe Virus Interferes with S-acylation of Remorin and Induces Its Autophagic Degradation to Facilitate Virus Infection. Molecular Plant 11:269–287.

Goodin, M.M., Zaitlin, D., Naidu, R.A., and Lommel, S.A. 2008. Nicotiana benthamiana: its history and future as a model for plant-pathogen interactions. Mol Plant Microbe Interact 21:1015–1026.

Gronnier, J., Crowet, J.-M., Habenstein, B., Nasir, M.N., Bayle, V., Hosy, E., Platre, M.P., Gouguet, P., Raffaele, S., Martinez, D., Grelard, A., Loquet, A., Simon-Plas, F., Gerbeau-Pissot, P., Der, C., Bayer, E.M., Jaillais, Y., Deleu, M., Germain, V., Lins, L., and Mongrand, S. 2017. Structural basis for plant plasma membrane protein dynamics and organization into functional nanodomains. eLife 6:e26404.

Hicks, S.W., and Galán, J.E. 2013. Exploitation of Eukaryotic Subcellular Targeting Mechanisms by Bacterial Effectors. Nature reviews. Microbiology 11:10.1038/nrmicro3009.

Hoefle, C., and Hückelhoven, R. 2008. Enemy at the gates: traffic at the plant cell pathogen interface. Cellular Microbiology 10:2400–2407.

Hurley, B., Lee, D., Mott, A., Wilton, M., Liu, J., Liu, Y.C., Angers, S., Coaker, G., Guttman, D.S., and Desveaux, D. 2014. The Pseudomonas syringae type III effector HopF2 suppresses Arabidopsis stomatal immunity. PLoS One 9:e114921.

Jarsch, I.K., and Ott, T. 2011. Perspectives on remorin proteins, membrane rafts, and their role during plant-microbe interactions. Mol Plant Microbe Interact 24:7–12.

Jarsch, I.K., Konrad, S.S., Stratil, T.F., Urbanus, S.L., Szymanski, W., Braun, P., Braun, K.H., and Ott, T. 2014. Plasma Membranes Are Subcompartmentalized into a Plethora of Coexisting and Diverse Microdomains in Arabidopsis and Nicotiana benthamiana. Plant Cell 26:1698–1711.

Jiang, S., Yao, J., Ma, K.W., Zhou, H., Song, J., He, S.Y., and Ma, W. 2013. Bacterial effector activates jasmonate signaling by directly targeting JAZ transcriptional repressors. PLoS Pathog 9:e1003715.

Jones, J.D., and Dangl, J.L. 2006. The plant immune system. Nature 444:323–329.

Karimi, M., Inzé, D., and Depicker, A. 2002. GATEWAY(TM) vectors for Agrobacterium-mediated plant transformation. Trends Plant Sci 7:193–195.

Khan, M., Seto, D., Subramaniam, R., and Desveaux, D. 2018a. Oh, the places they’ll go! A survey of phytopathogen effectors and their host targets. Plant J 93:651–663.

Khan, M., Youn, J.Y., Gingras, A.C., Subramaniam, R., and Desveaux, D. 2018b. In planta proximity dependent biotin identification (BioID). Sci Rep 8:9212.

Lal, N.K., Nagalakshmi, U., Hurlburt, N.K., Flores, R., Bak, A., Sone, P., Ma, X., Song, G., Walley, J., Shan, L., He, P., Casteel, C., Fisher, A.J., and Dinesh-Kumar, S.P. 2018. The Receptor-like Cytoplasmic Kinase BIK1 Localizes to the Nucleus and Regulates Defense Hormone Expression during Plant Innate Immunity. Cell Host & Microbe 23:485–497.e485.

Laluk, K., Luo, H., Chai, M., Dhawan, R., Lai, Z., and Mengiste, T. 2011. Biochemical and genetic requirements for function of the immune response regulator BOTRYTIS-INDUCED KINASE1 in plant growth, ethylene signaling, and PAMP-triggered immunity in Arabidopsis. Plant Cell 23:2831–2849.

Lee, A.H.-Y., Hurley, B., Felsensteiner, C., Yea, C., Ckurshumova, W., Bartetzko, V., Wang, P.W., Quach, V., Lewis, J.D., Liu, Y.C., Boernke, F., Angers, S., Wilde, A., Guttman, D.S., and Desveaux, D. 2012a. A Bacterial Acetyltransferase Destroys Plant Microtubule Networks and Blocks Secretion. PLoS Pathogens 8.

Lee, A.H., Hurley, B., Felsensteiner, C., Yea, C., Ckurshumova, W., Bartetzko, V., Wang, P.W., Quach, V., Lewis, J.D., Liu, Y.C., Börnke, F., Angers, S., Wilde, A., Guttman, D.S., and Desveaux, D. 2012b. A bacterial acetyltransferase destroys plant microtubule networks and blocks secretion. PLoS Pathog 8:e1002523.

Lefebvre, B., Timmers, T., Mbengue, M., Moreau, S., Herve, C., Toth, K., Bittencourt-Silvestre, J., Klaus, D., Deslandes, L., Godiard, L., Murray, J.D., Udvardi, M.K., Raffaele, S., Mongrand, S., Cullimore, J., Gamas, P., Niebel, A., and Ott, T. 2010. A remorin protein interacts with symbiotic receptors and regulates bacterial infection. Proc Natl Acad Sci U S A 107:2343– 2348.

Lewis, J.D., Wu, R., Guttman, D.S., and Desveaux, D. 2010. Allele-specific virulence attenuation of the Pseudomonas syringae HopZ1a type III effector via the Arabidopsis ZAR1 resistance protein. PLoS Genet 6:e1000894.

Lewis, J.D., Abada, W., Ma, W., Guttman, D.S., and Desveaux, D. 2008. The HopZ Family of Pseudomonas syringae Type III Effectors Require Myristoylation for Virulence and Avirulence Functions in Arabidopsis thaliana. J Bacteriol 190:2880–2891.

Lewis, J.D., Lee, A., Ma, W., Zhou, H., Guttman, D.S., and Desveaux, D. 2011. The YopJ superfamily in plant-associated bacteria. Mol Plant Pathol 12:928–937.

Lewis, J.D., Wilton, M., Mott, G.A., Lu, W., Hassan, J.A., Guttman, D.S., and Desveaux, D. 2014. Immunomodulation by the Pseudomonas syringae HopZ type III effector family in Arabidopsis. PLoS One 9:e116152.

Lewis, J.D., Lee, A.H., Hassan, J.A., Wan, J., Hurley, B., Jhingree, J.R., Wang, P.W., Lo, T., Youn, J.Y., Guttman, D.S., and Desveaux, D. 2013. The Arabidopsis ZED1 pseudokinase is required for ZAR1-mediated immunity induced by the Pseudomonas syringae type III effector HopZ1a. Proc Natl Acad Sci U S A 110:18722–18727.

Liang, P., Stratil, T.F., Popp, C., Marín, M., Folgmann, J., Mysore, K.S., Wen, J., and Ott, T. 2018. Symbiotic root infections in Medicago truncatula require remorin-mediated receptor stabilization in membrane nanodomains. Proceedings of the National Academy of Sciences 115:5289–5294.

Lin, W., Ma, X., Shan, L., and He, P. 2013. Big Roles of Small Kinases: The Complex Functions of Receptor-Like Cytoplasmic Kinases in Plant Immunity and Development. Journal of Integrative Plant Biology 55:1188–1197.

Liu, Y., Schiff, M., and Dinesh-Kumar, S.P. 2002a. Virus-induced gene silencing in tomato. Plant J 31:777–786.

Liu, Y., Schiff, M., Marathe, R., and Dinesh-Kumar, S.P. 2002b. Tobacco Rar1, EDS1 and NPR1/NIM1 like genes are required for N-mediated resistance to tobacco mosaic virus. Plant J 30:415– 429.

Lu, D., Wu, S., He, P., and Shan, L. 2010. Phosphorylation of receptor-like cytoplasmic kinases by bacterial Flagellin. Plant signaling & behavior 5:598–600.

Ma, W., Dong, F.F., Stavrinides, J., and Guttman, D.S. 2006. Type III effector diversification via both pathoadaptation and horizontal transfer in response to a coevolutionary arms race. PLoS Genet 2:e209.

Macho, A.P., and Zipfel, C. 2015. Targeting of plant pattern recognition receptor-triggered immunity by bacterial type-III secretion system effectors. Curr Opin Microbiol 23C:14-22.

Marín, M., and Ott, T. 2012. Phosphorylation of Intrinsically Disordered Regions in Remorin Proteins. Frontiers in plant science 3:86.

Marín, M., Thallmair, V., and Ott, T. 2012. The Intrinsically Disordered N-terminal Region of AtREM1.3 Remorin Protein Mediates Protein-Protein Interactions. The Journal of Biological Chemistry 287:39982–39991.

Nguyen, H.P., Chakravarthy, S., Velasquez, A.C., McLane, H.L., Zeng, L., Nakayashiki, H., Park, D.H., Collmer, A., and Martin, G.B. 2010. Methods to study PAMP-triggered immunity using tomato and Nicotiana benthamiana. Mol Plant Microbe Interact 23:991–999.

Perraki, A., Binaghi, M., Mecchia, M.A., Gronnier, J., German-Retana, S., Mongrand, S., Bayer, E., Zelada, A.M., and Germain, V. 2014. StRemorin1.3 hampers Potato virus X TGBp1 ability to increase plasmodesmata permeability, but does not interfere with its silencing suppressor activity. FEBS Lett 588:1699–1705.

Raffaele, S., Mongrand, S., Gamas, P., Niebel, A., and Ott, T. 2007. Genome-wide annotation of remorins, a plant-specific protein family: evolutionary and functional perspectives. Plant Physiol 145:593–600.

Raffaele, S., Bayer, E., Lafarge, D., Cluzet, S., German Retana, S., Boubekeur, T., Leborgne-Castel, N., Carde, J.P., Lherminier, J., Noirot, E., Satiat-Jeunemaitre, B., Laroche-Traineau, J., Moreau, P., Ott, T., Maule, A.J., Reymond, P., Simon-Plas, F., Farmer, E.E., Bessoule, J.J., and Mongrand, S. 2009. Remorin, a solanaceae protein resident in membrane rafts and plasmodesmata, impairs potato virus X movement. Plant Cell 21:1541–1555.

Rao, S., Zhou, Z., Miao, P., Bi, G., Hu, M., Wu, Y., Feng, F., Zhang, X., and Zhou, J.-M. 2018. Roles of Receptor-Like Cytoplasmic Kinase VII Members in Pattern-Triggered Immune Signaling. Plant Physiology 177:1679–1690.

Reymond, P., Kunz, B., Paul-Pletzer, K., Grimm, R., Eckerskorn, C., and Farmer, E.E. 1996. Cloning of a cDNA encoding a plasma membrane-associated, uronide binding phosphoprotein with physical properties similar to viral movement proteins. The Plant Cell 8:2265–2276.

Seto, D., Koulena, N., Lo, T., Menna, A., Guttman, D.S., and Desveaux, D. 2017. Expanded type III effector recognition by the ZAR1 NLR protein using ZED1-related kinases. Nat Plants 3:17027.

Shao, F., Golstein, C., Ade, J., Stoutemyer, M., Dixon, J.E., and Innes, R.W. 2003. Cleavage of Arabidopsis PBS1 by a bacterial type III effector. Science 301:1230–1233.

Simon-Plas, F., Perraki, A., Bayer, E., Gerbeau-Pissot, P., and Mongrand, S. 2011. An update on plant membrane rafts. Current Opinion in Plant Biology 14:642–649.

Simons, K., and Gerl, M.J. 2010. Revitalizing membrane rafts: new tools and insights. Nat Rev Mol Cell Bio 11:688.

Son, S., Oh, C.J., and An, C.S. 2014. Arabidopsis thaliana Remorins Interact with SnRK1 and Play a Role in Susceptibility to Beet Curly Top Virus and Beet Severe Curly Top Virus. The Plant Pathology Journal 30:269–278.

Stall, R.E., Bartz, J.A., and Cook, A.A. 1974. Decreased hypersensitivity to xanthomonads in pepper after inoculations with virulent cells of Xanthomonas vesictoria. Phytopathology 64:731–735.

Sun, J., Huang, G., Fan, F., Wang, S., Zhang, Y., Han, Y., Zou, Y., and Lu, D. 2017. Comparative study of Arabidopsis PBS1 and a wheat PBS1 homolog helps understand the mechanism of PBS1 functioning in innate immunity. Scientific Reports 7:5487.

Toth, K., Stratil, T.F., Madsen, E.B., Ye, J., Popp, C., Antolin-Llovera, M., Grossmann, C., Jensen, O.N., Schussler, A., Parniske, M., and Ott, T. 2012. Functional domain analysis of the Remorin protein LjSYMREM1 in Lotus japonicus. PLoS One 7:e30817.

Wang, G., Roux, B., Feng, F., Guy, E., Li, L., Li, N., Zhang, X., Lautier, M., Jardinaud, M.F., Chabannes, M., Arlat, M., Chen, S., He, C., Noel, L.D., and Zhou, J.M. 2015. The Decoy Substrate of a Pathogen Effector and a Pseudokinase Specify Pathogen-Induced Modified-Self Recognition and Immunity in Plants. Cell Host Microbe 18:285–295.

Witzel, K., Buhtz, A., and Grosch, R. 2017. Temporal impact of the vascular wilt pathogen Verticillium dahliae on tomato root proteome. J Proteomics 169:215–224.

Yamada, K., Yamaguchi, K., Shirakawa, T., Nakagami, H., Mine, A., Ishikawa, K., Fujiwara, M., Narusaka, M., Narusaka, Y., Ichimura, K., Kobayashi, Y., Matsui, H., Nomura, Y., Nomoto, M., Tada, Y., Fukao, Y., Fukamizo, T., Tsuda, K., Shirasu, K., Shibuya, N., and Kawasaki, T. 2016. The Arabidopsis CERK1-associated kinase PBL27 connects chitin perception to MAPK activation. The EMBO Journal 35:2468–2483.

Zhang, J., Li, W., Xiang, T., Liu, Z., Laluk, K., Ding, X., Zou, Y., Gao, M., Zhang, X., Chen, S., Mengiste, T., Zhang, Y., and Zhou, J.M. 2010. Receptor-like cytoplasmic kinases integrate signaling from multiple plant immune receptors and are targeted by a Pseudomonas syringae effector. Cell Host Microbe 7:290–301.

Zheng, X.Y., Spivey, N.W., Zeng, W., Liu, P.P., Fu, Z.Q., Klessig, D.F., He, S.Y., and Dong, X. 2012. Coronatine promotes Pseudomonas syringae virulence in plants by activating a signaling cascade that inhibits salicylic acid accumulation. Cell Host Microbe 11:587–596.

Zhou, H., Lin, J., Johnson, A., Morgan, R.L., Zhong, W., and Ma, W. 2011. Pseudomonas syringae type III effector HopZ1 targets a host enzyme to suppress isoflavone biosynthesis and promote infection in soybean. Cell Host Microbe 9:177–186.

